# A semi-supervised approach uncovers thousands of intragenic enhancers differentially activated in human cells

**DOI:** 10.1101/020362

**Authors:** Juan González-Vallinas, Amadis Pagés, Babita Singh, Eduardo Eyras

## Abstract

**Background:** Transcriptional enhancers are generally known to regulate gene transcription from afar. Their activation involves a series of changes in chromatin marks and recruitment of protein factors. These enhancers may also occur inside genes, but how many may be active in human cells and their effects on the regulation of the host gene remains unclear.

**Results:** We describe a novel semi-supervised method based on the relative enrichment of chromatin signals between 2 conditions to predict active enhancers. We applied this method to the tumoral K562 and the normal GM12878 cell lines to predict enhancers that are differentially active in one cell type. These predictions show enhancer-like properties according to positional distribution, correlation with gene expression and production of enhancer RNAs. Using this model, we predict 10,365 and 9,777 intragenic active enhancers in K562 and GM12878, respectively, and relate the differential activation of these enhancers to expression and splicing differences of the host genes.

**Conclusions:** We propose that the activation or silencing of intragenic transcriptional enhancers modulate the regulation of the host gene by means of a local change of the chromatin and the recruitment of enhancer-related factors that may interact with the RNA directly or through the interaction with RNA binding proteins. Predicted enhancers are available at http://regulatorygenomics.upf.edu/Projects/enhancers.html

## Background

Transcriptional enhancers are characterized by specific chromatin signatures, which differ depending of whether the enhancer is active or not [1,2,3,4,5]. Transcriptional enhancers have been generally characterized by studying the genome-wide binding of the acetyl-transferase P300, a ubiquitous enhancer co-activator [1,6,7]. However, not all P300-bound enhancers show activity [8]. Enhancers have also been characterized by their chromatin state [1,2,9,10]; and, although the mono-methylation of histone 3 lysine 4 (H3K4me1) has been identified to be an important signature for enhancers [2], this mark is not sufficient for enhancer activation [3,11]. In fact, recent evidence shows that other marks like H3K27ac [1,3,4,5] and H3K4me3 [5,11] may be necessary for enhancer activity. Additionally, the recruitment of RNAPII and the concomitant production of enhancer-associated RNAs (eRNAs) have also been associated to active enhancers [3,4,5,12,13].

Although enhancers are typically defined to regulate gene transcription at a distance, about 50% of potential enhancers predicted by high-throughput methods lie within protein-coding genes [2] and some overlap exons [14,15]. Intragenic enhancers can regulate the expression of the host gene [14] or of a nearby gene [15], and have been proposed to act as alternative promoters [16]. These results raise the question of how many intragenic enhancers may be active in a cell and whether upon their activation or silencing they may affect the processing of the host gene, possibly by means of local changes of the chromatin state. In this direction, there is evidence that some enhancers upstream of a reporter gene can affect splicing in vitro [17], and that intragenic enhancers bound by Argonaute-1 (AGO1) protein can affect the constitutive and alternative splicing of the host gene [18]. In this work we describe a computational method to predict active enhancers based on chromatin signals. This method, which uses the relative enrichment of chromatin signals between cell lines to the detect cell specific active enhancers, predicts thousands of intragenic active enhancers. Additionally, we find evidence that the differential activation of enhancers inside genes affect the expression and splicing of the host genes. We propose that the activation or silencing of intragenic transcriptional enhancers can modulate the regulation of the host gene through a local change of the chromatin.

## Methods

### Datasets

Annotated human enhancers with a mouse homologous enhancer that had been experimentally validated were downloaded from VISTA [19]. The gene set was obtained from the 7^th^ release of GENCODE, human assembly GRCh37 (hg19). ChIP-Seq and RNA-Seq datasets were downloaded from ENCODE [20] for K562 and GM12878 cells. The datasets used were: ChIP-Seq for CTCF, EZH2, P300, RNAPII, PU.1 (SPI1), STAT1, H3K9ac, H3K27ac, H3K4me1, H3K4me2, H3K4me3, H3K27me3, H3K36me3, H3K79me2, H4K20me1 and H2A.Z; one Control ChIP-Seq experiment and one input experiment; RNA-Seq for short (<200nt) and long (>200nt), polyA+ and polyA- RNAs from whole cell, nucleus and cytosol; and DNaseI data for the same cell lines. All datasets were downloaded in the form of mapped reads to the reference hg19 genome in BAM format.

### Relative enrichment calculation

We considered sliding windows of 1500nt along the entire genome, as suggested by the length distribution of experimentally validated enhancers [19,21] (Supplementary Figure 1), with a slide shift of 500nt, resulting in a total of 3,086,047 overlapping windows. In order to avoid mixing enhancer signal with genic and promoter signals, we discarded windows that were closer than 500nt to an annotated TSS. The same approach was applied to intergenic (Supplementary Figure 2A) and intragenic (Supplementary Figure 2B) regions. Although there are more intergenic windows (∼3·10^6^ vs ∼2.2·10^6^) in both cases the amount of windows with signal was similar (∼1.5 million windows), which were then kept for further processing. The relative enrichment of chromatin signals between 2 cell lines was calculated to predict active enhancers in K562 (relative increase of activation marks in K562 with respect to GM12878) and silent enhancers in K562 (relative decrease of activation marks in GM12878 with respect to K562, i.e. active in GM12878). Full quantile normalization for counts and GC content was applied using EDASeq [22]. GC content in each region was calculated as the proportion of G+C in the 1500nt window. After normalization, the z-score of the relative enrichment of each ChIP-Seq signals between K562 and GM12878 was calculated with Pyicoteo [23] using the *pyicoenrich* function (https://bitbucket.org/regulatorygenomicsupf/pyicoteo). A vector of z-scores per region was obtained, which we refer to as attributes, consisting of the 17 enrichment z-scores for the ChIP-Seq and Input datasets. A positive z-score in for a region indicates an increased in ChIP-Seq signal in K562 relative to GM12878 in that region, whereas a negative z-score indicates a decreased signal in K562 relative to GM12878; and z-scores close to zero indicate no significant differences between the cell lines. For all datasets, except for the ChIP-Seq with non-specific antibody and for the RNA-Seq datasets, we used replicates. The relative enrichments were calculated with respect to the distribution described by the comparison between replicates. When replicates were not available, these were simulated by pooling the two conditions and dividing them using random sampling [23].

**Figure 1.**
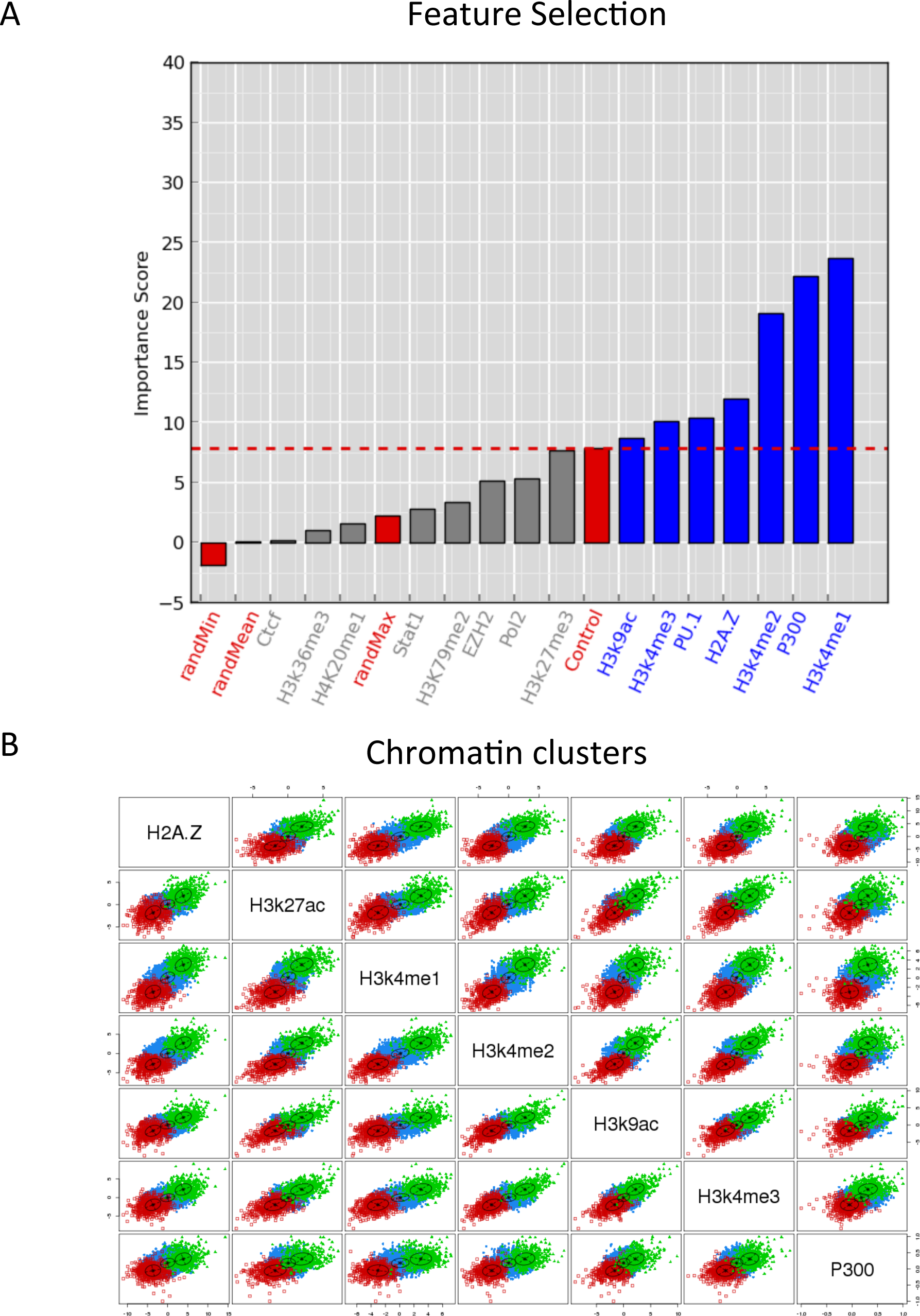
A predictive model of active enhancers. **(A)** Feature selection using H3K27ac (minus the Input DNA) as a correlation class. The bars represent the average importance score per feature. Red labels and bars indicate the minimum (randMin), mean (randMean) and maximum (randMax) of the simulated replicates, as well as the ChIPKSeq with a non-specific antibody (Control). The red dashed line separates the relevant features (in blue) from the non-relevant features (in grey). (**B)** ScaPer plot of the intergenic windows according to relative enrichment zKscores for every pair of selected features (x and y axis). Each dot represents a window and windows are separated according to the three classes: active enhancers (green), noKchange regions (blue) silent enhancers (red). The black centroids show the centers and standard deviations of the correlations between different features

**Figure 2.**
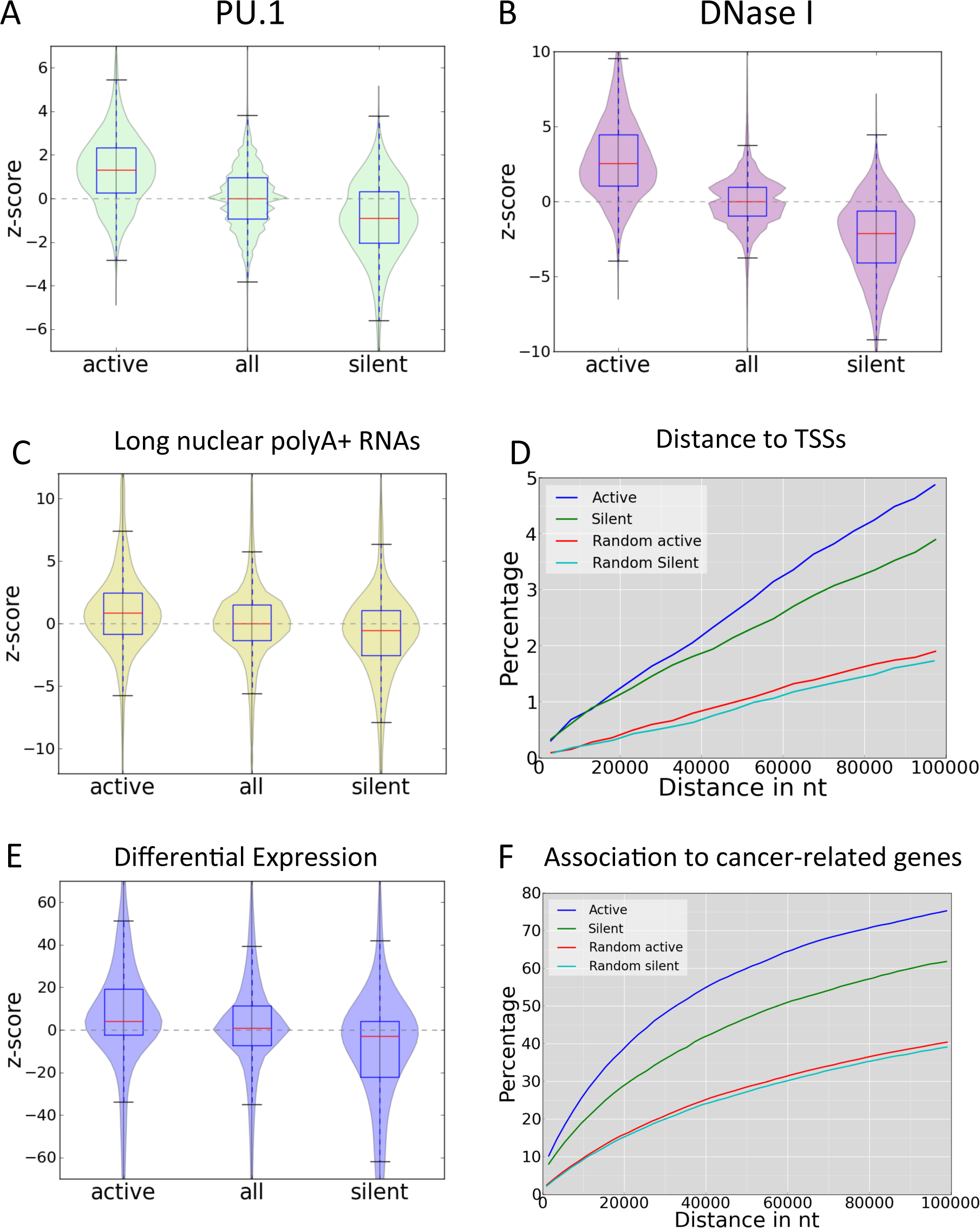
Properties of predicted intergenic enhancers. Relative enrichment of PU.1 **(A)**, DNaseI **(B)** and polyadenylated long (>200nt) nuclear RNA **(C)** at active and silent enhancer, as well as for regions of noKchange in chromatin. The violin plots describe the distributions for the zKscore of the relative enrichment along the yKaxis. Positive zKscore values mean enrichment in K562, while negative zKscores mean enrichment in GM12878. **(D)** Percentage of enhancers at a given distance from the TSS, for active (blue), silent (green), as well as for the corresponding randomized sets (red and cyan) (Methods). **(E)** Relative expression change in genes associated to enhancers by proximity to the TSS. The violin plot describes the distributions of zKscores of the relative enrichment of RPKM values along the yKaxis for genes associated to active and silent enhancers, as well as for noKchange regions calculated with Pyicoteo [23]. Genes where linked to the nearest enhancers within a distance range between 2kb and 10kb. **(F)** Cumulative distribution of enhancer nearby genes related to cancer in terms of the distance between the TSS and the closest enhancer. The comparison is made between active and silent predicted enhancers, and the corresponding randomizations.

### Feature Selection

Feature selection was performed using Boruta [24], which finds informative features by measuring the relevance of each attribute with respect to a reference attribute, also called correlation class, and in comparison with a random model extracted from the original dataset. Boruta uses the correlation class to evaluate the other features against it using Random Forests [25]. We performed this analysis using as correlation class each of the individual marks (Supplementary Figure 3). In each case, the correlation was performed 10 times using normalized counts on a subset of 5000 intergenic windows, sampled randomly in each one of the 10 iterations. In order to avoid possible biases, in each analysis the correlation class was defined as the ChIP-Seq signal minus the level of Input DNA. The features used as negative controls were the ChIP-Seq sample for a non-specific antibody (Control sample) (Supplementary Figure 3D) and H4K20me1 (Supplementary Figure 3E), which has been associated to transcription repression and heterochromatin but not to enhancer activity [26,27]. Running the selection algorithm with the H3K4me1 mark, the average Boruta score for the control increased notably, suggesting that the mark is present in many regions along the genome (Supplementary Figure 3C).

**Figure 3.**
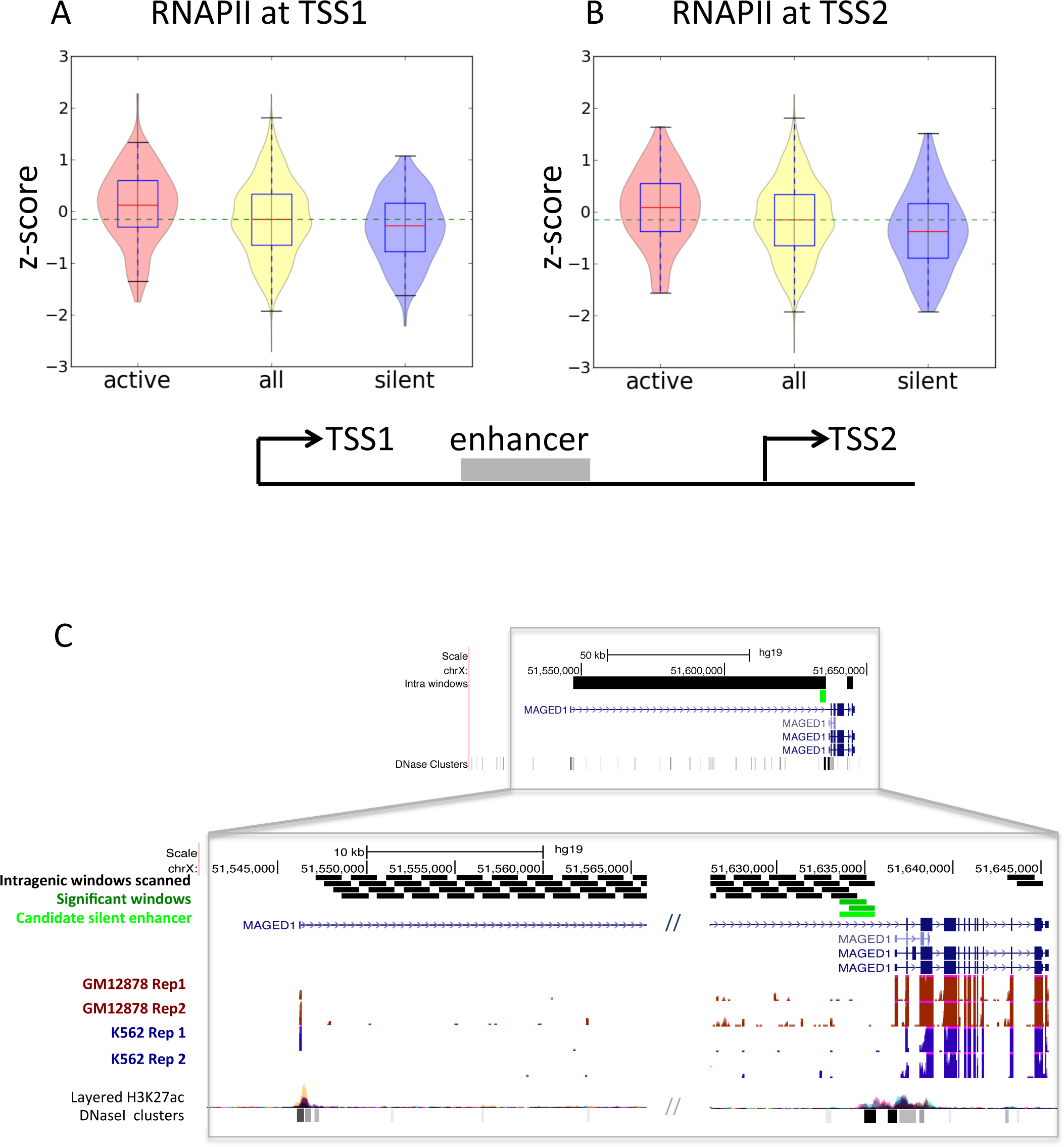
Intragenic enhancers and differential transcript expression. Relative enrichment zK scores of RNAPII on the most upstream TSS (TSS1) **(A)** and the second TSS (TSS2) **(B)** when there is an active (left red violin plots) or silent (right blue violin plots) enhancer between both TSSs and it sits at a minimum distance of 1000 nt from either one. The yellow violin plots in the middle represent the z-score distribution of all TSS1KTSS2 pairs, with or without predicted enhancers and the dashed green line represents the median value of this distribution. **(C)** A TSS1KTSS2 event in gene MAGED1 with a silent enhancer in K562 close to TSS2 (light green box). The black boxes indicate the intragenic 1500nt windows analyzed to search for enhancers. Dark green boxes indicate the significant windows and light green indicates the candidate silent enhancer. RNAKSeq data from K562 and GM12878 cell lines (2 replicates each) is shown below, together with the H3K27ac and DNase I hypersensitive sites tracks from ENCODE. The long intron separating the two TSSs is cut in the middle for simplicity.

### Window clustering

15000 arbitrary intergenic windows of length 1500bp were used as seed for the prediction model. Various different seed selections of the same size did not change the results significantly. These 15000 windows were clustered using Mclust [28]. Mclust is based on finite normal mixture modeling and uses the Bayesian Information Criterion (BIC) [29] for model optimization. The BIC score plateaus at 3 clusters for most models (Supplementary Figure 4A). The seed windows corresponded to 552 active, 616 silent and 13,832 no-change windows. This indicates that there are mostly three main classes, two that correspond to active and silent enhancers, and a class composed of a gradient of multiple chromatin states, which show little or no relative change of chromatin activity. This is further supported by the uncertainty plot, which shows that regions classified with higher certainty are on the extreme values of the correlation (Supplementary Figure 4B). The final model used for clustering was the centroid type (labeled as VEV), which creates clusters with variable volume, equal shape, and variable orientation. This model was used to classify the genome-wide 1500bp windows (Supplementary Figure 2) using the same clustering method Mclust to predict intergenic enhancers. Intragenic enhancers were calculated using the same seed of 15,000 intergenic windows as before. The clustering was performed in the same way as for intergenic enhancers. As controls we calculated 4 sets of randomized positions (intergenic/intragenic and active/silent putative predictions). These sets were calculated from the predicted enhancers by randomizing the positions, not closer than 500nt to any gene, avoiding gaps, genic regions, and other random locations previously generate, and keeping the same length and the same number of regions (Supplementary Figure 2A). Random intragenic enhancers were generated similarly by placing the intragenic enhancers in a random location inside the same gene, avoiding regions of 1kb around any internal TSSs and avoiding other random enhancers previously generated (Supplementary Figure 2B). All predicted intergenic and intragenic enhancers can be visualized in the UCSC genome browser through the link http://regulatorygenomics.upf.edu/Projects/enhancers.html

**Figure 4.**
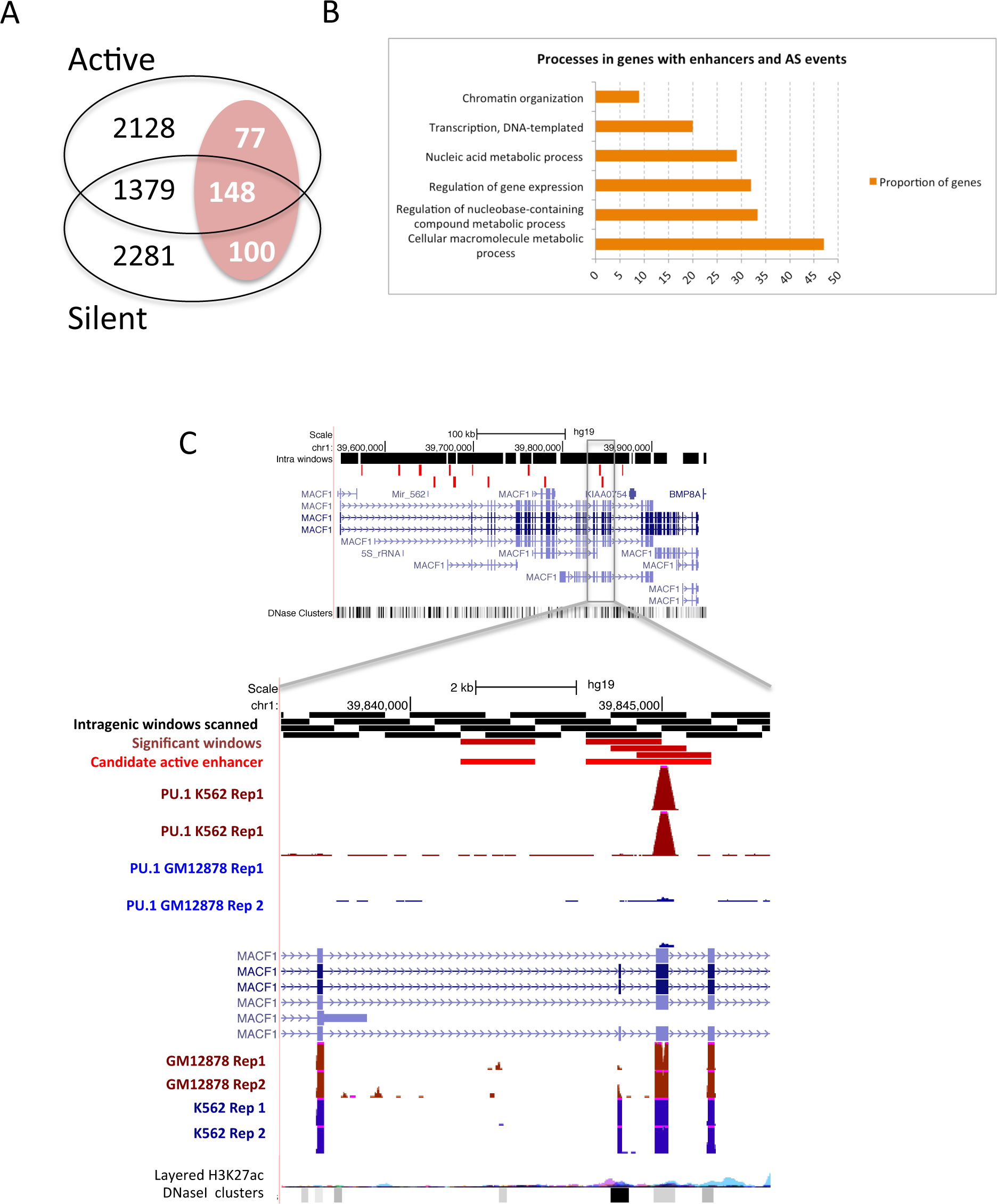
Effect of intragenic enhancers on splicing. **(A)** From the 2205 genes found to have active enhancers, 77 of them have regulated events. From the 1527 genes with active and silent enhancers, 148 have regulated events, and from 2381 genes with silent enhancers, 100 have regulated events. **(B)** Enriched Gene Ontology processes in the genes with active or silent enhancers and with regulated events, compared to genes with intragenic enhancers but no regulated events (Supplementary Table 3). **(C)** Example of a regulated alternative splicing event in the gene MACF1. The cassePe exon shows increased inclusion in K562 with deltaKPSI = 0.72 (PSI = 0.86 in K562 and PSI = 0.14 in GM12878). Two active enhancers are predicted in K562 in the area of the regulated exon (light red boxes). Intragenic scanned windows are indicated as black boxes, and significant windows are dark red boxes. One of the predicted enhancer regions shows binding of PU.1 in K562 (2 replicates) but no biding in GM12878 (2 replicates). RNAKSeq data from K562 and GM12878 cell lines (2 replicates each) is shown below, together with the H3K27ac and DNase I hypersensitive sites tracks from ENCODE.

### Linking enhancers to genes

Enhancers were linked to genes by selecting the closest TSS on either direction and by using ChIA-PET data for RNAPII in K562 cells for two replicates from ENCODE [20]. An enhancer was considered connected to a gene if there were at least 3 ChIA-PET pairs connecting both the predicted enhancer and the region of 1kb around the TSS of the gene. Random enhancers used as controls were calculated as described above. For the association of enhancers to genes, only enhancers that were between 2kb and 100kb from a TSS were considered. Genes associated to cancer were obtained from the Cancer Gene Census (http://cancer.sanger.ac.uk/cancergenome/projects/census/) [30].

### Expression and splicing analysis

For every gene in GENCODE (v7) annotation [31], the most upstream TSS (TSS1) and all alternative TSSs (TSS2, TSS3, etc.) were considered. Each pair TSS1-TSS2, TSS2-TSS3, etc. was considered as an alternative transcription event. RNAPII relative enrichment levels were measured around each TSS using the same method as before. To control possible association with upstream enhancers, we discarded all alternative TSS events that had a predicted intergenic enhancer (active or silent) 100kb upstream of the gene. We calculated the expression levels of the annotated transcript isoforms using cufflinks v2.1.1 [32] with parameters *--library-type fr-firststrand --no-effective-length-correction --min-frags-per-transfrag 5* and masking all rRNAs, tRNAs and mitochondrial sequences. The relative changes in transcript abundance were obtained using Cuffdiff with parameters *--library-type fr-firststrand --min-reps-for-js-test 1*, using the merged GTF file obtained from Cufflinks for GM12878 and K562, along with the bam files of GM12878 and K562 with replicates. This provided 3552 genes (6.68%) with relative changes in expression between the two cell lines.

Alternative splicing events from the Gencode v7 annotation [31] were calculated using the software SUPPA (https://bitbucket.org/regulatorygenomicsupf/suppa). Only events that do not overlap any other alternative splicing event were kept, giving rise to a total of 5319 events. For exon skipping events, defined by an exon triple E1 – E2 – E3, the inclusion level (PSI) of the middle exon E2, was calculated as the fraction of reads that include the exon over the total number of reads that include and skip the exon:

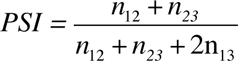

where n_12_, n_23_ and n_13_ are the number of reads that span the junctions E1-E2, E2-E3 and E1-E3, respectively. PSI values were calculated using junction reads only, since enhancers can produce RNA as well, so the enhancer-related RNAs may be mistakenly included in the PSI calculation when they overlap an event. Reads at junctions were counted with sjcount [33] from the mapped RNA-Seq data, using the *-read1 1* and *-read2 0* parameters. For this analysis, RNA-Seq reads were mapped using STAR [34] with parameters *-- outSJfilterOverhangMin -1 -1 -1 -1 and --sjdbScore 100* in order to use only annotated junctions. A genome index was previously generated with STAR over the Gencode.v7 annotation using the *--sjdbOverhang 75* parameter in order to adjust the splice junction database to the length of the RNA-Seq reads. Finally, only events with a total of 20 or more reads mapping at the junctions were kept. This gave a final number of 3227 and 3192 events with PSI values from the nuclear and cytosolic RNA-Seq experiments, respectively.

We defined the events to be regulated if they had |delta PSI| > 0.1 in at least one replicate comparison between cell lines. Using two pairings of the replicates, this gave rise to 339 and 293 events (148 in common) with the cytosolic samples, and 367 and 378 (210 in common) for the nuclear samples. Additionally, we defined a set of alternative events that do not change splicing by imposing |delta PSI| < 0.05 between the same replicate comparisons used before. This gave rise to 1722 and 1534 (1328 in common) for the cytosolic samples, and 1627 and 1497 (1278 in common) for the nuclear samples.

## Results and Discussion

### Modeling and prediction of active transcriptional enhancers

We built a computational predictive model based on the relative differences in various chromatin marks between two cellular conditions. We applied this model to study the differences between the ENCODE cell lines K562, a leukemia cell line, and GM12878, a blood cell derived cell line. Using windows along the entire genome (Supplementary Figures 1 and 2), we considered the relative enrichment of a number of histone marks and protein factors (Methods). We then clustered windows into classes according to the chromatin features. In order to determine which features are relevant for classification, we performed a feature selection analysis in which one signal is chosen as a proxy for a classification value and is compared against the rest (Methods). We then considered two of the main epigenetic marks related to active enhancers, H3K27ac [3] and H3K4me3 [11], as proxies for enhancer activity. We found that H3K4me1 and H3K4me2, observed to be present in active and non-active enhancers [11] are strongly correlating signals (Figure 1A and Supplementary Figure 3A), We also consistently found H2A.Z, which is a histone variant associated to open chromatin and H3K4 methylation [35]; and P300, which is ubiquitously present in enhancers [6].

Interestingly, when P300 or H3K4me1 were used as a correlation feature, the signals H3K27ac and H3K4me3 did not appear as the most significantly associated (Supplementary Figures 3B and 3C). Additionally, P300 seemed to associate with the largest subset of features, which is consistent with experimental evidence showing that P300 associates generally to enhancers [1,6]. However, enhancers with H3K4me1 and/or P300 occupancy are not always active [3,11], since H3K4me1 precedes enhancer-binding factors and P300 may be present in poised and intermediate enhancer states [36]. On the other hand, we did not find RNAPII and H3K36me3 to be strong predictors of enhancer activity (Figure 1A and Supplementary Figure 3A), even though they have previously detected on enhancers [12,13]. Additionally, although we found a strong correlation of PU.1 (SPI1) with H3K27ac, it does not correlate with H3K4me3, hence it is likely that PU.1 associates to a subset of the putative enhancers [37]. Based on these results, we decided to keep those features that scored consistently above the technical and biological controls in the feature selection analysis using H3K27ac and H3K4me3 as correlation classes, including these two marks. That is, we used as predictors of enhancer activity the following signals: P300, H3K27ac, H3K9ac, H3k4me1/me2/me3 and H2A.Z.

Clustering the genomic windows according to the relative enrichment of the selected features (Methods) resulted in three optimal classes (Supp. Figure 4). We recovered one class characterized for being enriched in H3K4me3 and H3K27ac (Figure 1B), which we considered to be enhancers that are active in K562 cells (silent in GM12878). We recovered a second class characterized by a depletion of these same marks in K562 (Figure 1B), which we considered to be active enhancers in GM12878 (silent in K562). Finally, the third cluster showed small or no changes in most of the signals, indicating that these regions do not have any differential activity between the two cell lines. These regions do not necessarily represent enhancers and are labeled as no-change. These three groups (active, silent, no-change) define the three predictable classes of our computational model, two of which can be identified with enhancer classes: active and silent. The genome wide classification analysis resulted in 66,079 windows predicted to be active in K562 (silent in GM12878) and 64,436 windows predicted to be active in GM12878 (silent in K562).

### In-silico validation of active transcriptional enhancers

In order to evaluate the accuracy of our predictions, we first compared our predicted enhancers windows with the enhancer regions predicted in the same cell lines by ChromHMM [38]. The majority of our enhancers predicted as active in K562 or GM12878 overlap with ChromHMM windows labeled as weak or strong enhancers in the same cells (Supplementary Figure 5A and 5B). On the other hand, when we compared active windows with ChromHMM labels in the other cell line, the majority corresponds to ChromHMM silent windows (Supplementary Figures 5C and 5D), as expected. Furthermore, the overlap of our active enhancers with predicted ChromHMM enhancers increases with the posterior probability of our predictions (Supplementary Figures 5E and 5F). In contrast, when comparing the active enhancers in one cell line with the ChromHMM labels from the other cell line, we found no correlation with the posterior probability (Supplementary Figures 5G and 5H). Based on these comparisons, we kept predictions with a posterior probability of > 0.95, which resulted in 36,301 active windows in K562 and 37,859 active windows in GM12878. Overlapping windows were then clustered into 16,646 active enhancers in K562 and 16,328 active enhancers in GM12878, which distribute evenly along the genome (Supplementary Figure 6A). These enhancers have mean length of 3,053bp and the majority of them (87.65%) are shorter than 5kb (Supplementary Figure 6B). There were also 273 (1.38%) predictions longer than 10kb, which may correspond to large-scale chromatin domains [39] or to clusters of enhancers [40]. We filtered out those predictions longer than 5kb, resulting in 10,365 active enhancers and 9,777 silenced enhancers, with mean lengths of 2,704.6bp and 2,588bp, (median lengths of 2,500bp and 2,000bp), respectively. These average lengths are in agreement with previous analyses of enhancers from ChIP-Seq data of histone marks and protein factors [5,11,15].

We next studied the association of enhancers to other signals not considered in the predictive model. PU.1 and RNAPII correlate with the predicted active enhancers, with 25.3% and 20.1% of the active enhancers in K562 showing a significant relative enrichment (left-tailed p-value < 0.01) in PU.1 (Figure 2A) and RNAPII (Supplementary Figure 7A), respectively. Similarly, we found a strong association of DNaseI to our predicted enhancers, in agreement with previous observations [2,9,10] and 53.6% of the active and silent enhancers show a significant enrichment (left-tailed p-value < 0.01) (Figure 2B). Likewise, 46.7% of the silent enhancers in K562 (active in GM12878) show a significant depletion in DNaseI (right-tailed p-value < 0.01). In contrast, H3K27me3 shows a weak inverse correlation with enhancer activity and 6.5% of the silent enhancers in K562 show a significant enrichment (right-tailed p-value < 0.01) of H3K27me3 (Supplementary Figure 7B). Although CTCF and H3K36me3 have been detected before on enhancers [12,13], we observed a weak correlation of these signals with enhancer activity and only 7.4% and 4.6% of active enhancers in K562 show a significant enrichment in CTCF and H3K36me3, respectively (Supplementary Figures 7C and 7D).

We additionally investigated whether enhancer-associated RNAs (eRNAs) are found in our predictions. Enhancer activity correlates with the production of polyA+ (Figure 2C) and polyA-(Supplementary Figure 8A) long (>200bp) nuclear RNAs, compared to silent enhancers. This relative enrichment is much larger than for the other RNA subclasses (Supplementary Figure 8B). Interestingly, there is also enrichment of cytosolic polyA+ RNAs (Supplementary Figure 8C), but not of cytosolic polyA- RNAs (Supplementary Figure 8D) or short RNAs (<200bp) (Supplementary Figure 8E and 8F). Moreover, not all enhancers predicted as active appear to generate eRNAs: 26.4% and 32.1% of the predicted active enhancers in K562 have a significant (left-tailed p-value < 0.01) increase of nuclear polyA+ and polyA-, respectively. In comparison, only 1.25% of active enhancers have significant (left-tailed p-value < 0.01) increase for short nuclear RNAs. For cytosolic polyA+, 18.7% of the predicted active enhancers in K562 have a significant (left-tailed p-value < 0.01) increase of eRNAs. In contrast, only 5% and 9.2% of these active enhancers have a significant enrichment of short total RNAs and polyA- cytosolic RNAs, respectively.

Although enhancers can regulate genes from afar, they tend to be enriched upstream of genes (Visel et al. 2009). We therefore connected enhancers to genes by choosing for each enhancer the closest annotated transcription start site (TSS) in either direction. With this approximation, active intergenic enhancers show enrichment at distances close to TSSs compared to random regions and to silent enhancers (Figure 2D). Using these enhancer-TSS pairs, we calculated the relative change in gene expression measured from RNA-Seq data (Methods). We observed that genes with activated enhancers at a distance between 2kb and 10kb show up-regulation, whereas genes with silenced enhancers in the same distance range show down-regulation (Figure 2E). Moreover, this association is conserved when the distance range of the enhancers is extended to be between 10kb and 100kb from the closest gene (Supplementary Figure 9A). Further support for transcription activity in association to our predicted enhancers was found measuring the relative density of RNAPII around the TSS in genes close to predicted enhancers, which was found to correlate with enhancer activity (Supplementary Figure 9B).

We additionally searched for evidence of direct physical interactions for the enhancer-TSS pairs calculated above by using ChIA-PET data for RNAPII [41]. Although only a small fraction of active enhancers have ChIA-PET links to TSS regions (1.6%), there is enrichment over silent enhancers and randomized regions (Supplementary Figure 9C), indicating that predicted active enhancers tend to have more ChIA-PET links than silent enhancers and expected by chance. Finally, we investigated whether enhancers active in K562 have any association to genes that have been involved in cancer. Using the cancer gene census [30], we found that enhancers predicted to be active in K562 are enriched for genes related to cancer, compared to random regions and to enhancers silent in K562 (active in GM12878) (Figure 2F). Interestingly, oncogenes can be linked more frequently to active enhancers and suppressors can be linked more frequently to silent enhancers (Supplementary Figure 10). In summary, these analyses indicate that our predicted enhancers show properties of active enhancers. We therefore set out to predict intragenic enhancers using the same computational model.

### Thousands of intragenic enhancers are differentially activated in human cells

Active enhancers regulating the expression of nearby genes have been observed in exons [14,15] and about 50% of enhancers predicted by high-throughput methods lie within protein-coding genes [2]. Additionally, by comparing the overlap of validated VISTA elements with the annotation in Gencode.v7 [31], we observe that there is no preference for intragenic or intergenic regions (Supplementary Figure 1). All these evidences indicate that intragenic enhancers represent an important regulatory component of the genome. However, it remains an open question how many intragenic enhancers may be active in a given cell. Accordingly, we decided to apply our predictive model to localize putative intragenic enhancers that are activated in K562 relative to GM12878, and vice versa.

In order to predict intragenic active enhancers, we considered 1.5kb sliding windows inside genes, starting 500bp downstream of the first TSS and eliminating all windows that overlap with a 1kb region around every annotated alternative TSS (Supplementary Figure 2). This resulted in an initial set of 2,206,307 possible 1.5kb windows, for which we used the same selected chromatin features as for the intergenic enhancers. Using a seed of 15000 intergenic regions and the same clustering approach as before, we predicted 73,080 active and 92,225 silenced regions. As we did previously with intergenic enhancers, we compared our predicted intragenic predictions with ChromHMM predictions with similar results (Supplementary Figure 11). Accordingly, we only kept windows predicted with posterior probability > 0.95, resulting in 42,297 and 55,624 active intragenic enhancer windows in K562 and GM12878, respectively. After clustering overlapping windows, we obtained 17,791 active intragenic enhancers in K562 (relative to GM12878) and 21,108 active intragenic enhancers in GM12878 (relative to K562), falling inside a total of 5,162 genes (10.11% of all genes) and 5,933 (11.61%) genes, respectively. The mean length of these predictions is 3,665bp, with the majority (82.81%) being shorter than 5kb (Supplementary Figure 12). As before, we kept those shorter than 5kb, resulting in 11,055 and 11,917 candidate active intragenic enhancers in K562 and GM12878, respectively.

Our predicted intragenic enhancers tend to occur in separate genes, with only 29.2% of the genes hosting enhancers of both types. The majority of intragenic enhancers active in K562 (78.24%) or active in GM12878 (80.61%) fall in intronic regions, and 26.02% in K562 (22.07% in Gm12878) overlap at least partially with an exon. However, comparing the proportion of exonic and intronic regions covered by enhancers with the actual proportions of these regions in genes, we find no preference for exons or introns (Supplementary Methods). Additionally, even though we observed a preference for intragenic enhancers to fall on the first intron (Supplementary Figure 13), this effect can be explained by the fact that first introns are on average longer in human (Supplementary Methods) [42].

### Intragenic enhancers affect the regulation of the host gene

As the activation or silencing of intragenic transcriptional enhancers are characterized by the differential density of chromatin marks, we hypothesized that this would lead to a modulation of the RNA processing of the host gene. To test this, we first measured whether genes hosting predicted enhancers tend to show significant differential expression between the two cell lines. Similarly as before for enhancers linked to genes, we find a correlation of the relative expression change of genes hosting active or silenced enhancers. Specifically, 23.8% of 5,162 genes with only active enhancers in K562 (34.5% of the 5,933 genes with only active enhancers in GM12878) show a significant expression up-regulation in the corresponding cell line (Methods). We then tested whether the activation or silencing of internal enhancers may produce the activation or repression of an intragenic TSS. We considered all active and silenced enhancers that fall between the most upstream TSS (TSS1) and the first internal annotated TSS (TSS2), such that the distance TSS1-TSS2 was longer than 20kb. This resulted in a total of 870 TSS1-TSS2 pairs, from which 113 (13%) had at least one active enhancer in K562 and 135 (15.52%) had at least one silent enhancer in K562 (active in GM12878) located between both TSSs. When an active enhancer is present between the two alternative TSSs, we observed that generally both TSS1 and TSS2 show an increase in RNAPII density in K562 relative to GM12878 (Figure 3A). This suggests that activation of an intragenic enhancer can affect both TSSs. Conversely, when a silent enhancer is present between both TSSs, the relative level of RNAPII tend to decrease at both TSSs relative to the other cell line (Figure 3B). Interestingly, this effect persists for other downstream alternative TSS events (Supplementary Figure 14), indicating that intragenic enhancers can activate internal TSSs, but also affect transcription of the most upstream TSS to some extent. We further used ChIA-PET for RNAPII in K562 to validate a possible direct interaction between our intragenic enhancers and the first TSS of each gene. Similarly as before, we observe a higher density of ChIA-PET links for active enhancers than for silent and random ones (Supplementary Figure 15). In this case 54.19% of the active intragenic enhancers have ChIA-PET links, compared to 36.73% in silent, 23.5% in random active and 17.5% in random silent (Supplementary Figure 15). This enrichment with respect to intergenic enhancers could be due to a higher density of RNAPII sites in intragenic regions. As an example of the described mechanism, we show the example of the gene MAGED1, a member of the melanoma antigen family D, which is known to have tumor-suppressor properties [43]. We predict an enhancer that is silent in K562 and active in GM12878, and is located between a distant TSS and an alternative downstream TSS (Figure 3C). The activation of this enhancer co-occurs with the expression of the downstream first exon in GM12878 cells, whereas the silencing of the enhancer co-occurs with lack of expression of this exon in K562 cells (Figure 3C). The RNA-Seq data suggests that the activation of this enhancer affects more strongly the usage of the TSS that is downstream.

The change in chromatin state induced by the activation or silencing of an enhancer may affect the processing of the pre-mRNA. There is evidence that localized intragenic chromatin states can produce changes in alternative splicing through various mechanisms [44,45,46,47,48]. Moreover, we have recently shown that active enhancers recruit Argonaute proteins to regulate the splicing of the host gene [18]. We therefore hypothesized that intragenic enhancers that are active in a cell line relative to the other one may in general be associated with relative differences in the inclusion level of nearby exons relative to the two same cell lines. To test this, we measured for all genes the variation in splicing between K562 and GM12878 using cytosolic and nuclear RNA-Seq polyA+ data from ENCODE, using only junction-reads, to avoid contributions from RNAs stemming from overlapping enhancers (Methods). We found that around 4% of multi-exonic genes with intragenic enhancers that are active in either cell line, have a regulated alternative splicing event (|delta PSI|>0.1) between both cell lines, whereas only about 1% of all the genes without intragenic enhancers have a calculated alternative splicing event that changes between cell lines (Supplementary Table 2). A total of 3732 and 3908 genes with no change in expression have active or silent enhancers, respectively, and 1527 of these genes have enhancers of both types (Figure 4A). Moreover, 325 of the genes with no change in expression have regulated events, and overlap with a total of 1046 enhancers (480 active and 566 silent) (Figure 4A). Moreover, these genes contain 347 of the 535 (65%) cassette events regulated between K562 and GM12878 (available as Supplementary File). Using Gorilla [49], we tested whether genes with enhancers and regulated events were enriched for any particular Gene Ontology term, and found an overrepresentation of genes encoding DNA-binding proteins implicated in gene regulation and chromatin organization (Figure 4B) (Supplementary Table 3).

We next decided to evaluate whether there is any association between the presence of enhancers and regulated events in genes. To this end, we compared only genes that have one or more of the 5319 calculated alternative splicing events (Methods) and separated these genes according to whether they have one or more regulated events (|delta PSI| > 0.1) or or not (|delta PSI| < 0.05) between the two cell lines. We found that in all comparisons the proportion of genes with regulated events was higher in those genes that have active enhancers (either in K562 or GM12878) (Supplementary Table 4), being the comparison statistically significant (Fisher p-value < 0.05) for both replicates for genes with active enhancers in GM12878, using nuclear RNA-Seq for the calculation of PSI values; whereas the same association for enhancers active in K562 was only significant for one of the replicate comparisons (Fisher p-value = 0.01) (Supplementary Table 4). Moreover, these associations remained significant when we considering only those genes that do not change expression between both cell lines (Supplementary Table 5). The regulated events in genes with active or silent intragenic enhancers present equal proportions of each pattern of PSI change, i.e. increase or decrease PSI (Supplementary Figure 16), which is consistent with the observed dual effect that a chromatin change can have on splicing [50]. Additionally, the direction of change of PSI does not correlate with the position, upstream or downstream, of the enhancer relative to the regulated exon (Supplementary Figure 17). Remarkably, the majority of the regulated events are located 5000nt from an enhancer (Supplementary Figure 18). However, we did not find any significant difference with the distribution of distances of non-regulated events to nearby enhancers (Supplementary Figure 19).

As an example, we show the case of a regulated exon in the microtubule-actin crosslinking factor 1 gene (MACF1) (Figure 4C). We observe a cassette exon with increased inclusion (delta PSI = 0.72) in K562 cells. The regulated exon is flanked by two enhancers predicted to be active in K562, one of which shows binding of PU.1 in K562, but not in GM12878 (Figure 4C). This, together with the rest of our findings, suggests that the binding of PU.1 to a nearby enhancer, possibly in combination with other factors, could control the inclusion of this exon in MACF1. MACF1 has been implicated in the Wnt signalling pathway [51] and the inclusion of a cassette exon in MACF1 was observed before to be associated to lung adenocarcinoma [52]. This result suggests the interesting possibility that the binding of PU.1 to an enhancer inside the MACF1 gene may affect its splicing, thereby altering Wnt signaling and contributing to the oncogenic transformation associated to PU.1 [53]. In conclusion, we have found a possible association between the activity of intragenic enhancers and the regulation of the pre-mRNA. In particular, we find evidence that the activation of intragenic enhancers, besides affecting the activity of internal TSSs, can also potentially influence the inclusion of nearby exons.

## Conclusions

We have developed a novel semi-supervised method that exploits the relative enrichment and depletion of multiple signals from ChIP-Seq experiments to predict enhancers that are active in one cell line relative to another. Applying this method to ENCODE data we predicted a total of 21,420 enhancers that are active in K562 relative to GM12878 (silent in GM12878 cells) and 21,694 enhancers that are active GM12878 relative to K562 (silent in K562), including intragenic and intergenic enhancers.

The number of active enhancers is cell type specific and very much dependent on the method used to detect them [36]. Although activation of enhancers is generally associated to a number of histone modifications, only a small fraction of the many candidate enhancers previously identified using a variety of techniques may be active in a given cell. For instance, Heintzman et al. found 24,566 putative enhancers in K562 cells with approximately 20% of them overlapping putative enhancers detected in HeLa cells [1]. In contrast, ChromHMM [38] predicts more than 60,000 non-abutting genomic regions to be strong enhancers and about three times as many for weak enhancers. There are two main reasons for the discrepancies with our predicted number of active enhancers: the resolution of the genome segmentation is very different and we only predict enhancers that are active in one condition but not in the other. That is, we do not detect enhancers active or silent in both cell types. Nonetheless, we found a good agreement between the regions we predict as active or silent enhancers and the annotations from ChromHMM for the same cell lines.

Our predicted enhancers are H3K27ac dependent and are defined almost entirely by chromatin signals. The relevant predictive features confirm that active enhancers are characterized not only by the presence of H3K4me1, but also by the presence of H3K27ac, H3K4me3 and RNAPII [4,5,12,13]. We also observed that active enhancers show an enrichment of the histone variant H2A.Z, which has been identified to demarcate regulatory regions [35]. In contrast, CTCF and EZH2 and the histone marks H3K36me3 and H4K20me1 do not seem to play any prominent role in enhancer activation. H3K27me3 is the only feature that shows a pattern of depletion in active enhancers and enrichment in silent enhancers, but mainly in long enhancer-like regions (data not shown), which may be related to other regulatory mechanisms. We additionally found that predicted enhancer activity correlates strongly with production of long nuclear RNAs, rather than short ones, which can be polyA+ as well as polyA-. However, we observe that not all active enhancers produce eRNAs. Furthermore, although RNAPII and H3K36me3 have been detected on enhancers in relation to eRNA production [12,13], we did not find them as strong predictors of enhancer activity.

When we applied the same predictive model to predict intragenic enhancers, we found a similar number of active intragenic enhancers as for intergenic ones. This result suggests that there exist in cells a considerable number of differentially activated intragenic enhancers, which may have a relevant contribution to the mechanisms of cell-specific gene regulation. Since active enhancers are characterized by a local modification of the chromatin state, we hypothesized that our predictions could be linked to relative differences between the same two cell lines in expression and splicing. We observed that intragenic transcriptional enhancers, upon activation or silencing, affect the activity of downstream alternative transcription start sites. Surprisingly, they can also affect the most upstream TSS. This generalizes previous findings indicating that intragenic enhancers can act as internal alternative promoters [16]. We also found that intragenic enhancers, upon activation or silencing, associate to the differential inclusion of nearby exons. However, a considerable proportion of splicing changes occur in genes that change expression (51.1% for genes containing differentially included exons in K562 and 52.9% for differentially excluded exons in GM12878). This indicates that the main effect of the activation of enhancers may be related to the activation of alternative transcription in the gene and alternative splicing may be a byproduct of that. The observed changes may be mediated by the changes in the RNAPII elongation produced due to the chromatin change. However, active intragenic enhancers show enrichment in open chromatin marks (H3K4me3, H3K27ac) that have not been associated before to changes in RNAPII elongation.

On the other hand, we found here a strong association of PU.1 (SPI1) to active enhancers in K562 cells and in particular, a significant increase in PU.1 occupancy in 26.8% of active enhancers. PU.1 has been shown before to be an essential co-factor for enhancer activity [54] and is known to bind to H3K4me1 sites in macrophages and B cells in a cell-specific manner [56,57]. Moreover, PU.1 has been observed to regulate alternative splicing from the promoter [58] and can interact with the RNA binding proteins FUS (TLS) and NONO (p54nrb) [59,60]. In fact, PU.1 has been proposed to bind RNA [60] and to perform an antagonistic function to the RNA binding proteins TLS and NONO in the regulation of splicing [60,61]. In this direction, we found enrichment of regulated events in genes with enhancers, which suggests that PU.1 could be regulating the splicing of some of these genes through its binding to intragenic enhancers, possibly interacting with RNA binding proteins [46]. In support of this model, we find that there is an enrichment of regulated events in genes with enhancers that are active or silent relative to the other cell line. We postulate that intragenic enhancers provide localized and cell-type specific mechanisms to link the chromatin state to RNA processing.

In summary, there is increasing evidence that changes in the chromatin state can affect the processing of the pre-mRNA [44,45,46,47,48,62,63,64,65,66] and different models for this regulation have been proposed. From our analysis a picture emerges whereby localised chromatin changes inside genes can take place by means of the activation of intragenic transcriptional enhancers. We propose that the differential activation and silencing of transcriptional enhancers that fall within genes could explain the localized chromatin variation that have been observed before to affect the expression and splicing of genes, either through the modulation of RNAPII activity or through the recruitment of factors that can interfere with RNA processing, like PU.1.

## Author contributions

EE designed and supervised the study, JGV carried out the enhancer analyses, AP and BS performed the splicing and expression studies, respectively. EE and JGV wrote the manuscript. All authors read and approved the final manuscript.

## List of abbreviations

ChIP: Chromatin Immunoprecipitation; Seq: sequencing; CTCF: CCCTC-binding factor; RNAPII: RNA Polymerase II; H3K36me3: Histone 3 Lysine 36 tri-methylation; H3K9me2: Histone 3 Lysine 9 de-methylation; H3K27me3: Histone 3 Lysine 27 tri-methylation; H3K27Ac: Histone 3 Lysine 27 acetylation; H3K4me1/2/3: Histone 3 Lysine 4 mono/di/tri methylation; H3K79me3: Histone 3 Lysine 79 tri-methylation; H4K20me1: Histone 4 Lysine 20 mono-methylation; H3K9Ac: Histone 3 Lysine 9 acetylation; H2A.Z: H2A Histone family member Z; STAT1: signal transducer and activator of transcription 1; polyA: poly-adenylation; BIC: Bayesian Information Criterion; TSS: transcription start site; ChIA-PET: chromatin interaction analysis by paired-end tag sequencing; PSI: percent spliced-in;

## Acknowledgements

The authors would like to thank E. Furlong, Y. Barash, B. Blencowe and U. Braunschweig for useful discussions. This work was supported by grants from Plan Nacional I+D (BIO2011- 23920) and Consolider (CSD2009-00080) from MINECO (Spanish Government), and by the Sandra Ibarra Foundation for Cancer (FSI 2013). JGV and BS were supported FPI grants from the MINECO (Spanish Government) BES-2009-018064 and BES-2012-052683, respectively.

## Supplementary Material

**Supplementary Figure 1.**
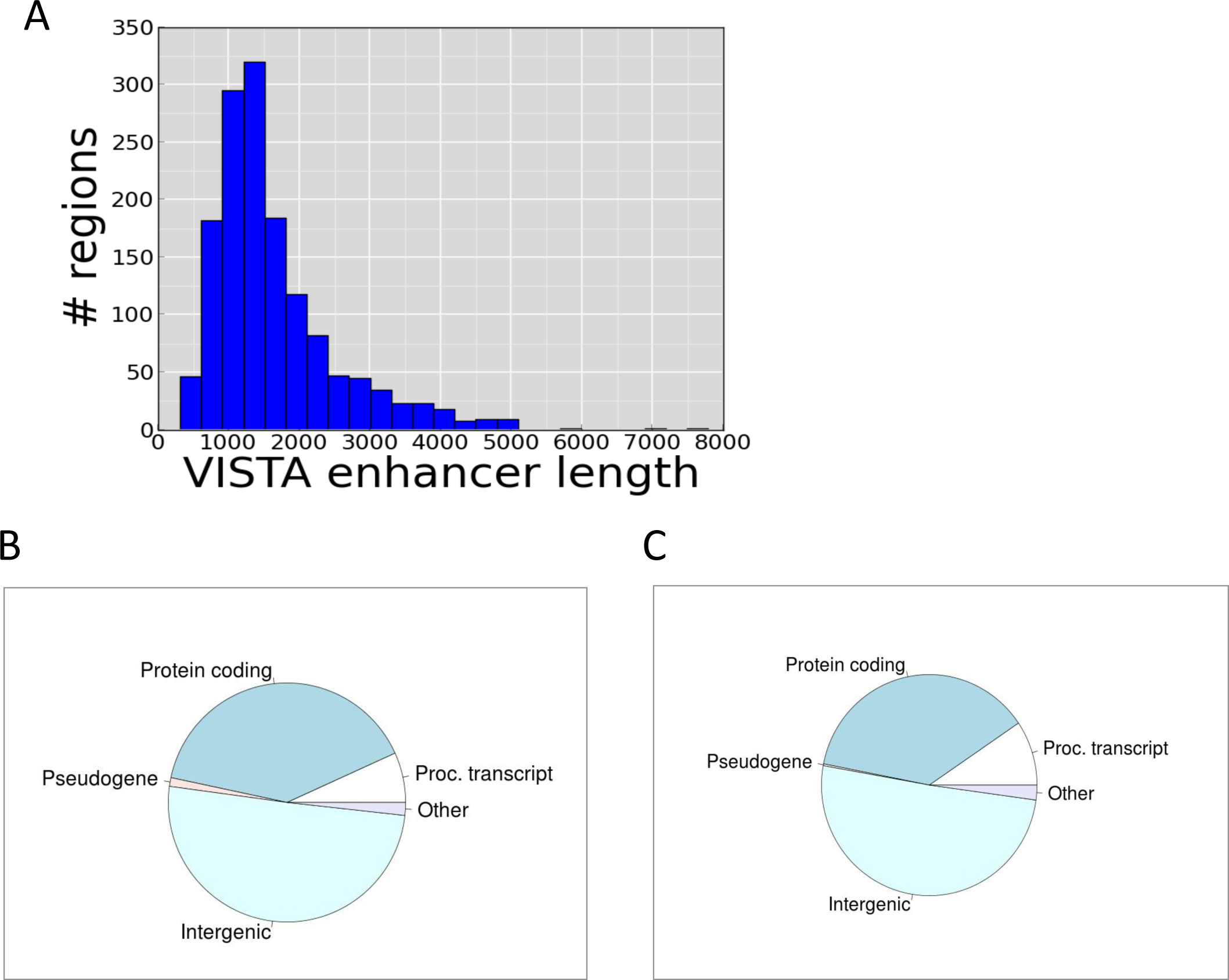
Properties of the VISTA enhancers. **A** - Length distribution of human VISTA enhancers. The average length of the VISTA regions is 1637.9 nt (median 1383nt), standard deviation 891nt. Out of the 1447 experimentally validated regions, only 6 (0.41%) are above 5000 nucleotides. **B** - GENCODE coverage of the genome. Pie chart representing the percentage of bases covered by different annotated elements in the human genome (GENCODE V7): protein coding genes, Processed Transcripts, Pseudogenes, regions and Other. The label Other summarizes Immunoglobulin (Ig) variable chain and TPcell receptor (TcR) genes (active and silent), several types of small non-coding RNA and lincRNA. The rest of non annotated elements by GENCODE are classiffed as intergenic. **C -** VISTA elements positioning. Percentage of the VISTA regions that fall in one of the categories described above.

**Supplementary Figure 2.**
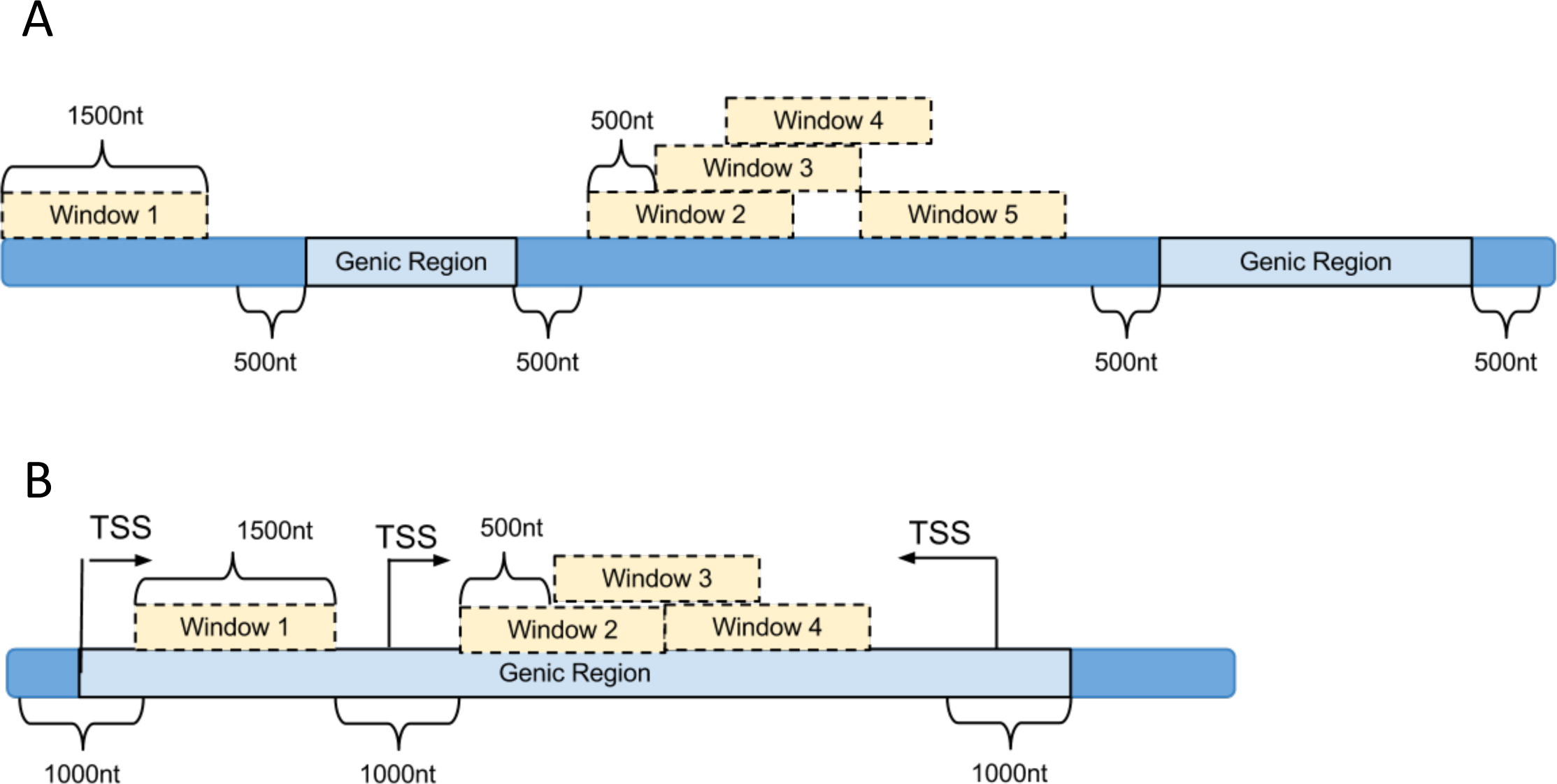
Intergenic and Intragenic exploratory windows. **A** - The intergenic exploratory intergenic windows have a size of 1500nt and are overlapping 500nt from each other. In order to avoid mixing with promoter signal, they are at least 500nt away from Genic regions. **B -** Intragenic windows have the same size as the intergenic windows and are at least 500nt away from any TSS (either first or intragenic).

**Supplementary Figure 3.**
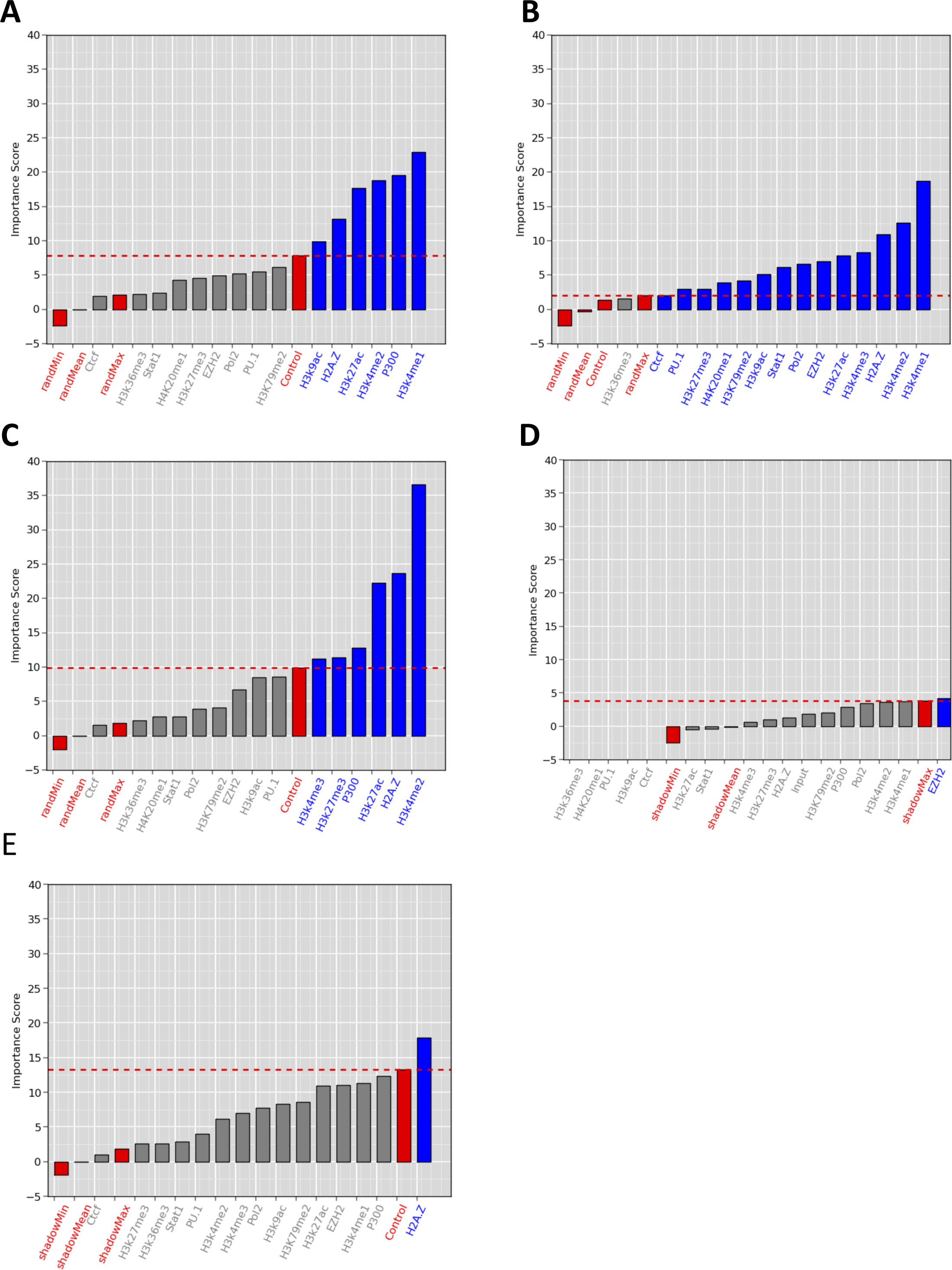
Feature Selection. **A** - Feature selection using H3K27ac as a correlation class. The bars represent the average importance score per feature after averaging over 10 random samples of 5000 intergenic windows extracting from all intergenic windows with non-zero signal in at least one cell type. Red labels and bars indicate the minimum (randMin), mean (randMean) and maximum (randMax) of the simulated replicates, as well as the ChIPPSeq with a non-specific antibody (Control). The red dashed line separates the relevant features (in blue) from the non-relevant features (in grey). **B** P Feature selection average scores using P300 as the correlation class. **C -** Feature selection average scores using H3K4me1 as the correlation class. **D -** Feature selection average scores using Control as the correlation class. The Control (ChIPPSeq experiment with no specific antibody) did not correlate significantly with any of the other features. On the other hand, **E -** Feature selection average scores using H4k20me1 as the correlation class. H4K20me1, which has been associated to transcription repression and heterochromatin (Balakrishnan et al. 2010, Beck et al. 2012) but not to enhancer activity, shows some correlation with H2A.Z, but no correlation with any other signal. Feature selection was performed using Boruta.

**Supplementary Figure 4.**
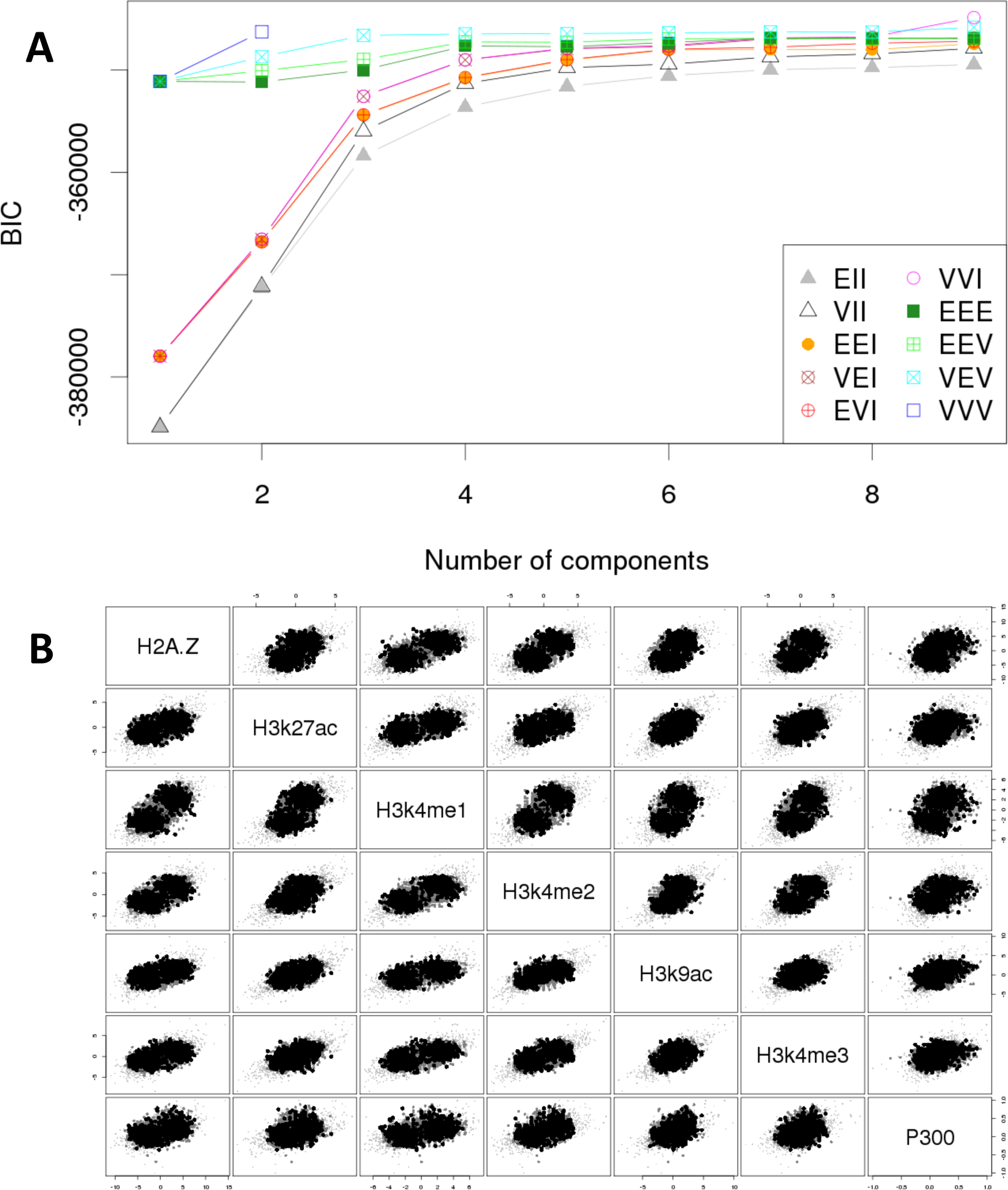
Intergenic Clustering. **A.** The Bayesian Information Criterion (BIC) calculation with Mclust. The X axis represents the number of clusters, the Y axis the BIC score. Every line correspond to a different type of model. The model that scores higher and plateaus faster is VEV (Variable Volume, Equal Shape, Variable Orientation). **B.** Mclust visual representation of the uncertainty. Dark dots represent more uncertainty (less probable to be good predictions), lighter dots are more certain (more probable to be good predictions). As expected, bigger differences between K562 and GM12878 levels correlate with less uncertainty.

**Supplementary Figure 5.**
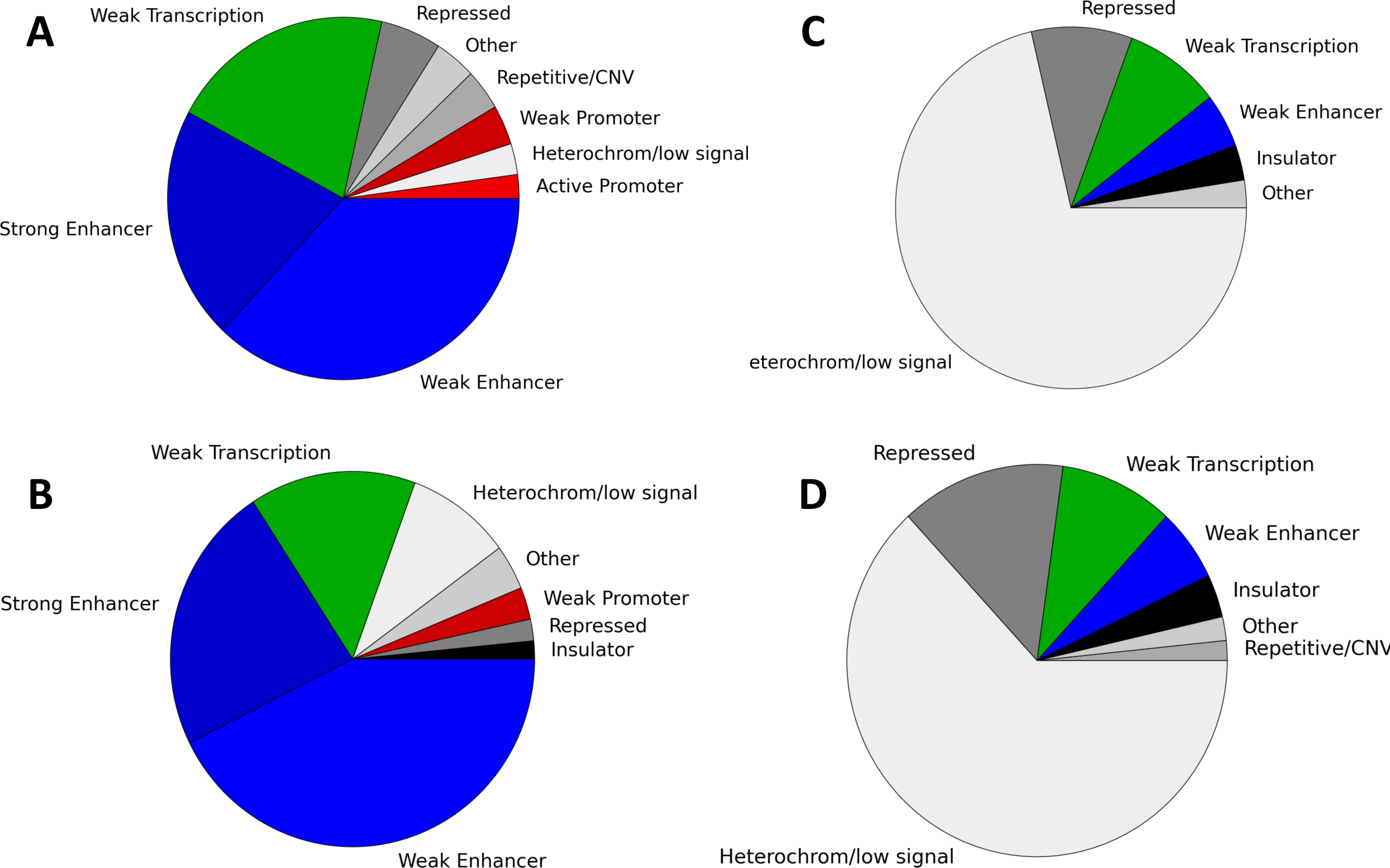
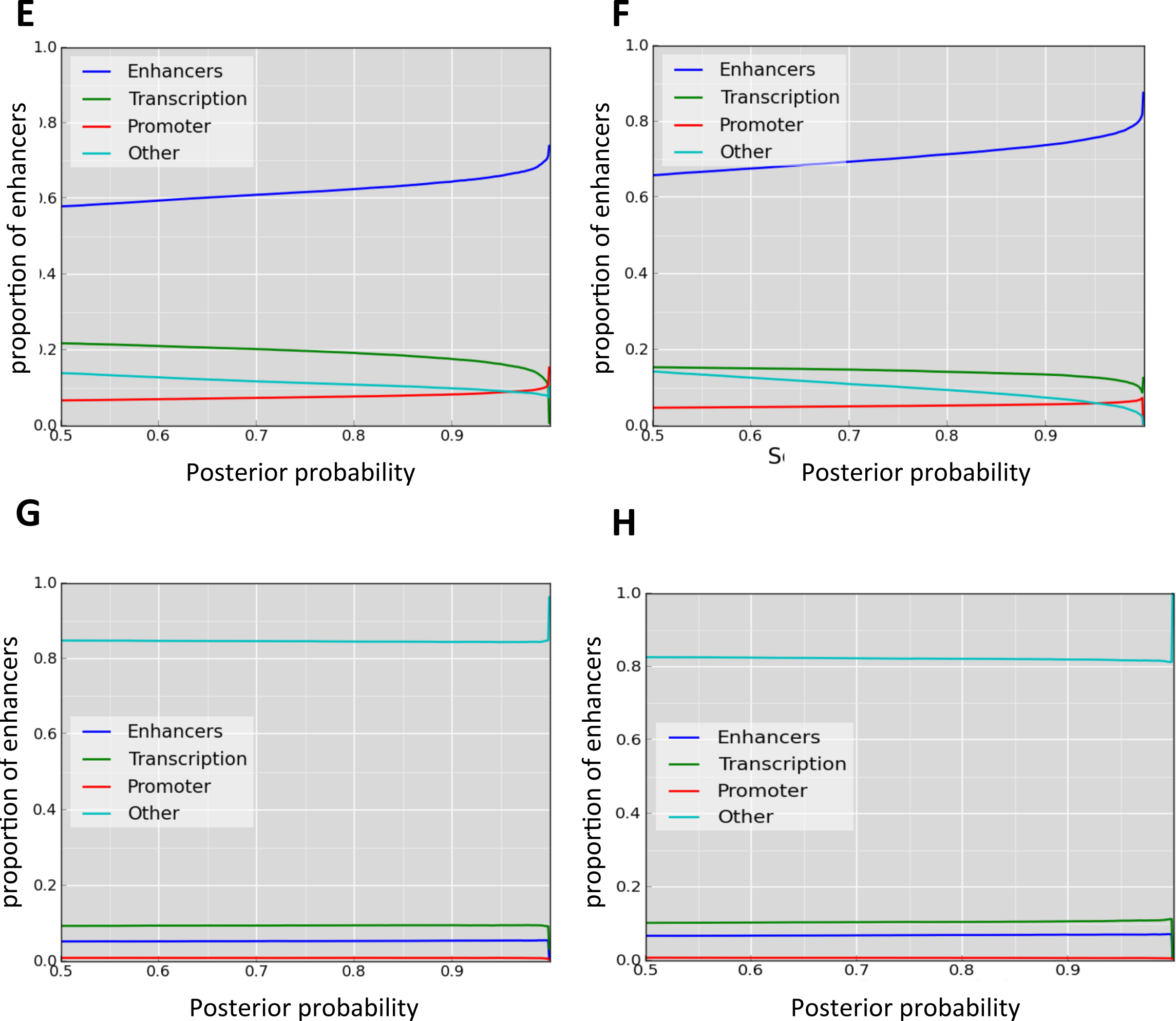
Comparing intergenic predictions with ChromHMM predictions. **A.** Pie chart showing the overlap of our intergenic windows predicted to be active in K562 with different ChromHMM (Ernst et al. 2011) classes for K562. Our predictions overlap with ChromHMM weak (light blue) or strong (dark blue) enhancers, weak transcription (green), weak (dark read) and active (light red) candidate promoters, repressed regions (dark grey), repetitive regions (middle grey) and others (light grey). **B.** As in A, pie chart showing the overlap of enhancers predicted to be active in GM12878 with the ChromHMM classes for GM12878. **C.** Pie chart showing the overlap of our windows predicted as active in K562 (silent in GM12878) with ChromHMM classes for GM12878. The majority of windows overlap with heterochromatin regions (light grey) and repressed regions (dark grey). There are also some overlaps with weak enhancers (blue) and weak transcription (green). **D.** As per C., Pie chart showing the overlap of our windows predicted as active intergenic enhancers in GM12878 (silent in K562) with ChromHMM classes for K562. The majority of windows overlap with heterochromatin regions (light grey) and repressed regions (dark grey). There are also some overlaps with weak enhancers (blue) and weak transcription (green). Proportion of predicted enhancers that are labeled by ChromHMM in each cell line (y axis) as a function of the posterior probability (x axi), for active enhancers in K562 **(E)** and GM12878 **(F)** and of silent enhancers in K562 **(G)** and GM12878 **(H).** The lines indicate the proportion of overlap ChromHMM regions labelled as weak or strong enhancers (blue), transcription related classes (green), promoters (red) and repressed (repressed, heterochromatin and repetitive) (cyan) regions

**Supplementary Figure 6.**
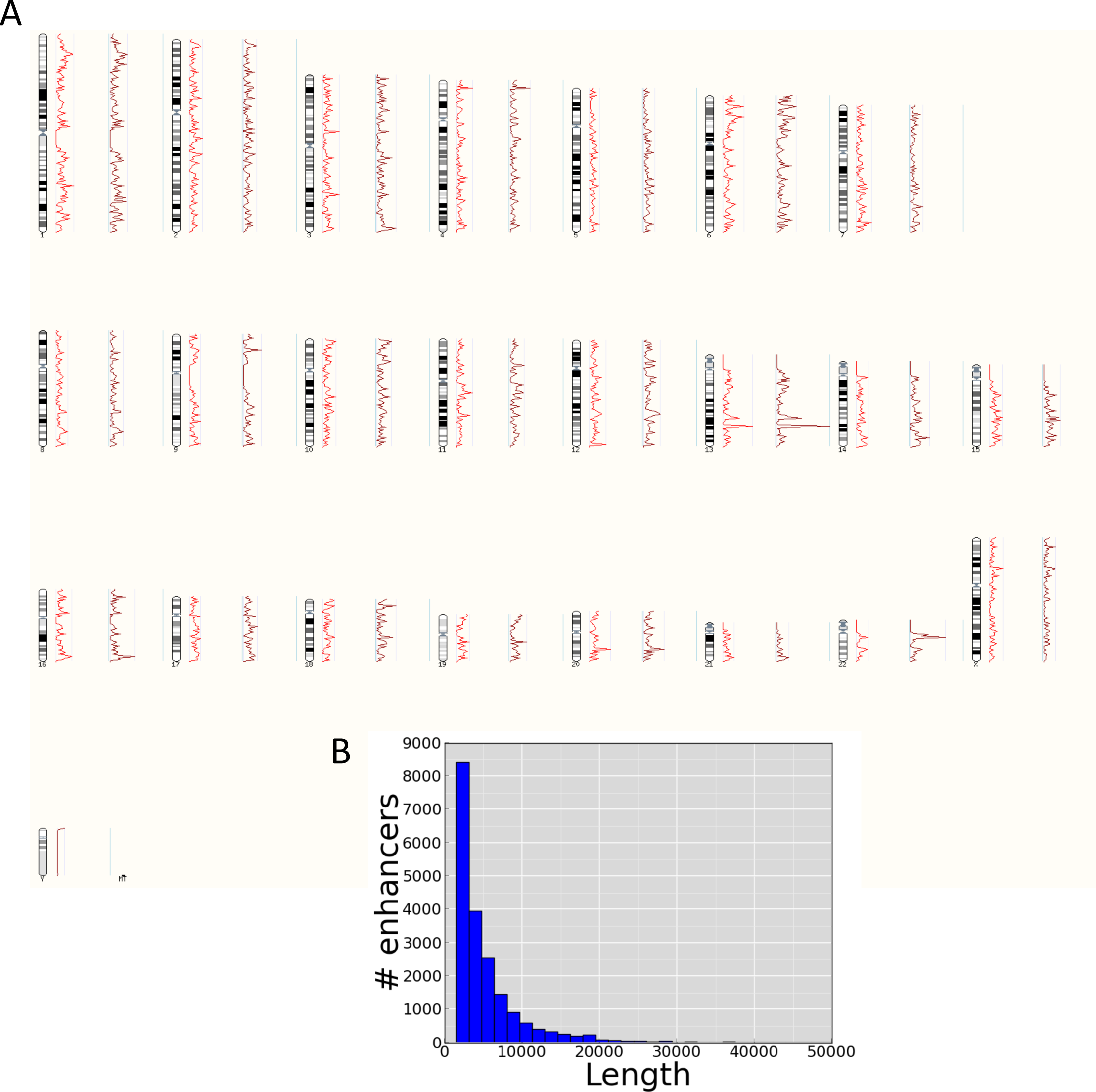
A. Genome wide distribution of intergenic enhancers. Profile of the density of predicted intergenic enhancers (active and silent) along the human karyotype, for all predicted enhancers (dark red) and for enhancers of lengths < 5kb (light red). **B -** Length distribution of our predicted intergenic enhancers

**Supplementary Figure 7.**
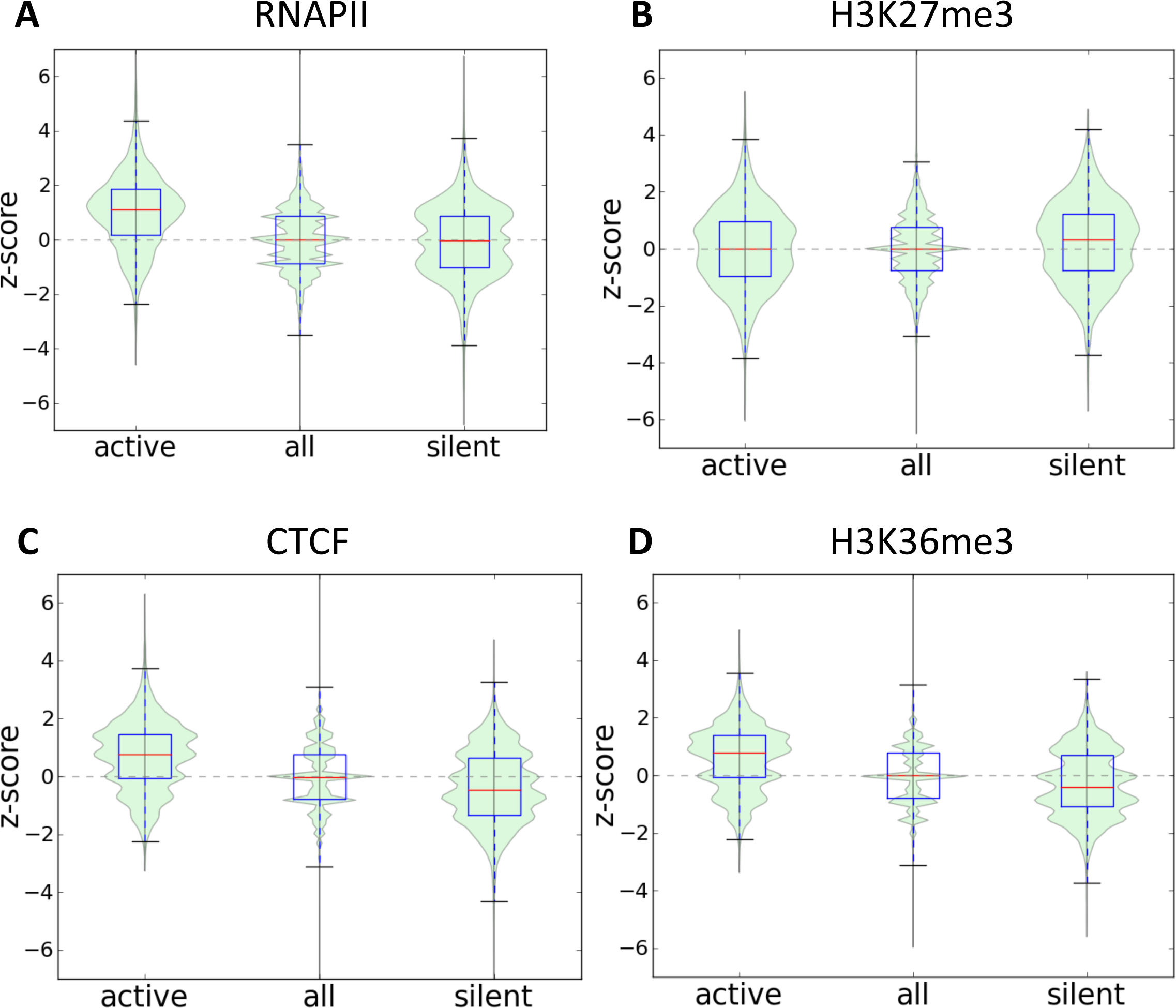
Relative enrichment of RNAPII **(A)**, H3K27me3 **(B)**, CTCF **(C)** and H3K36me3 **(D)** in intergenic enhancers. Z-score distributions for our putative active and silent enhancers, as well as for all regions

**Supplementary Figure 8.**
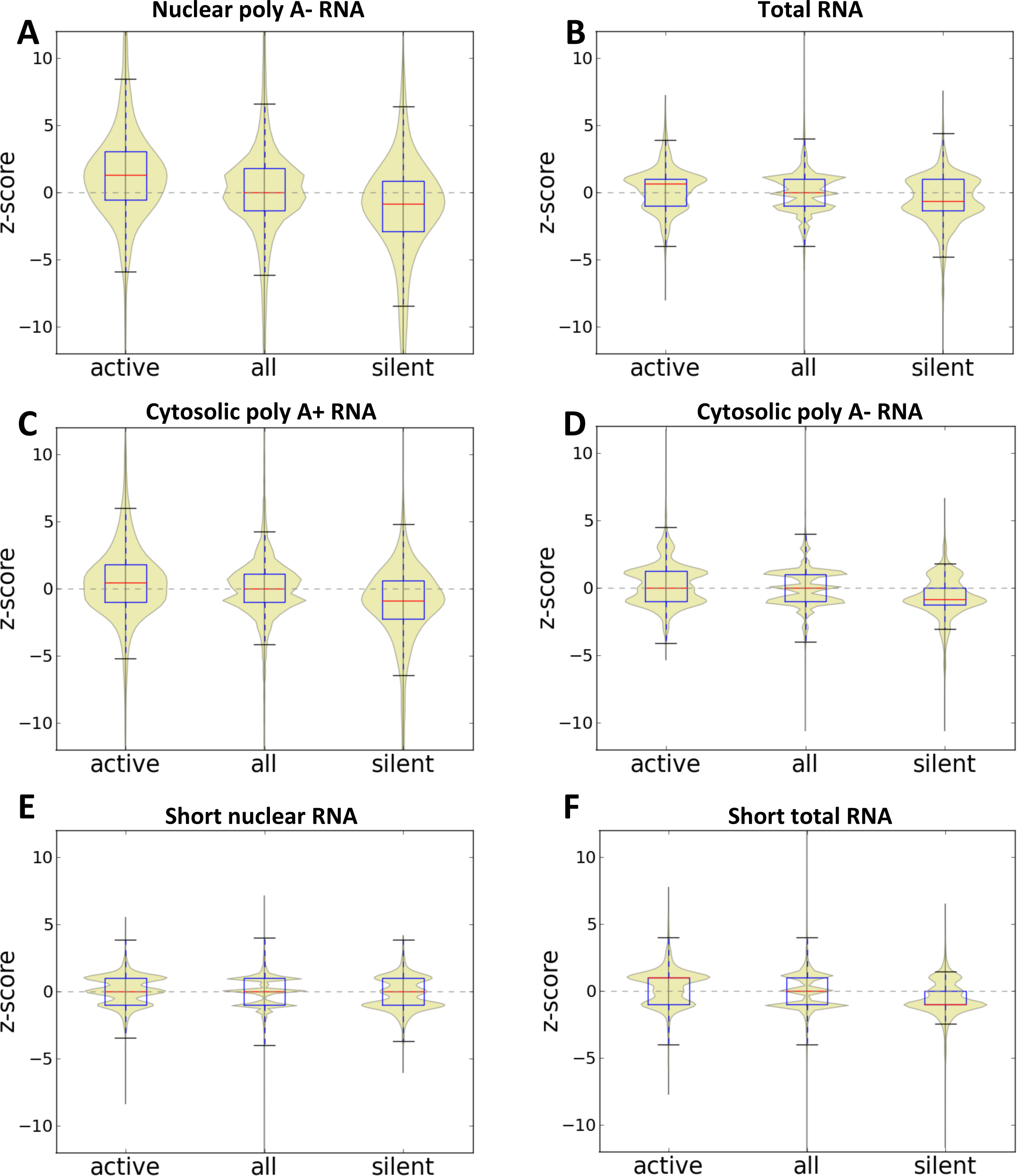
Violin plots for the relative enrichment in K562 relative GM12878 in active and silent enhancers, as well as the distribution of all zPscores, for nuclear long (>200nt) poly AP RNAs, **(B)** total RNA, **(C)** Cytosolic polyA+ RNAs, **(D)** Cytosolic polyAP RNAs, **(E)** Short (<200nt) nuclear RNAs and **(F)** Short (<200nt) Total RNAs.

**Supplementary Figure 9.**
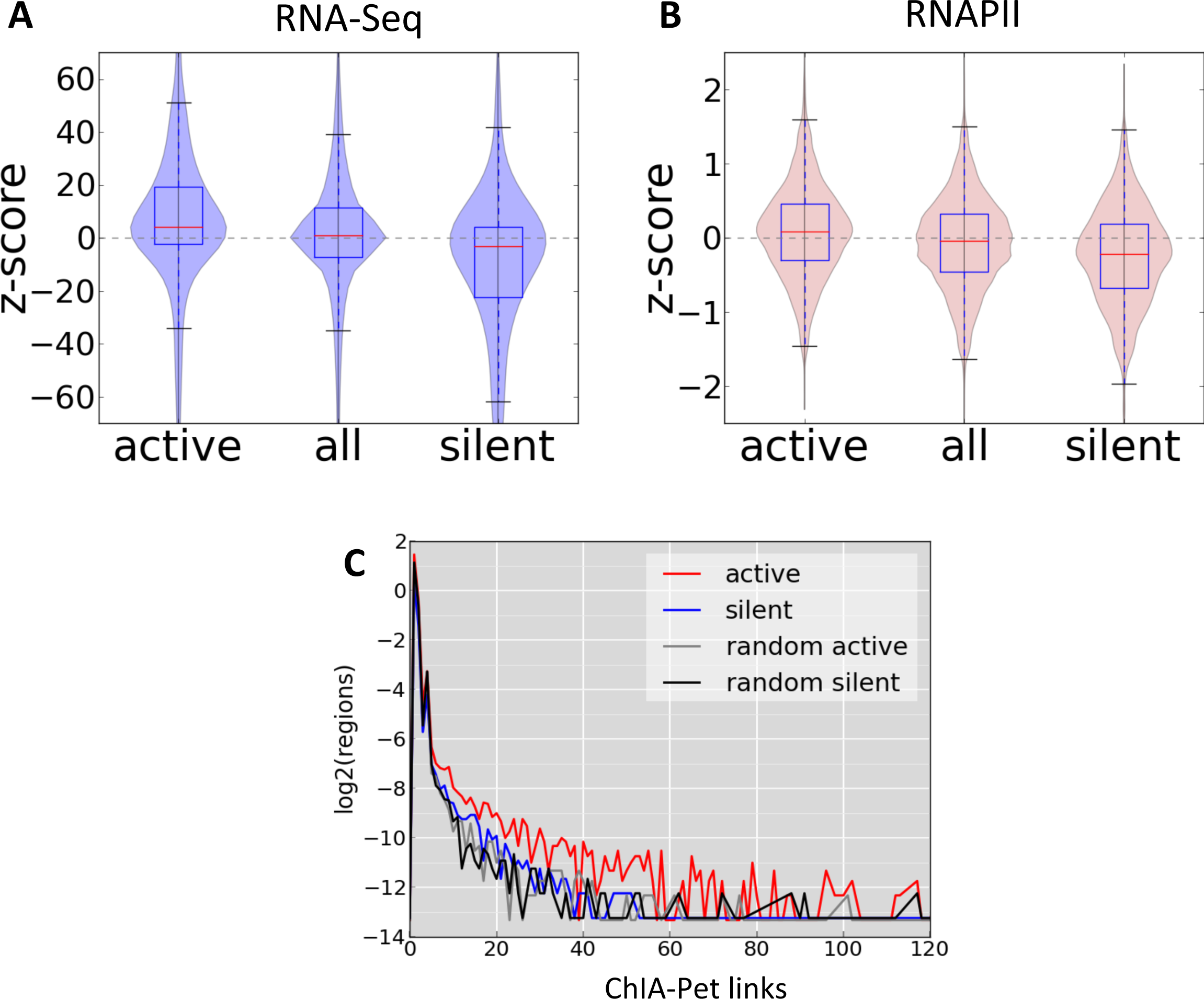
Relative change in **(A)** gene expression and in **(B)** RNAPII at the TSS for genes. The Y axis measures the relative expression change in terms of a ZPscore for genes associated to active (left violin plot) and silent enhancers (right violin plot), and for all genes (middle violin plots). The relative change in RNAPII density is measured in a 1kb window around the TSS. Genes were associated to the nearest predicted enhancer within a range of 10kb to 100kb from the TSS on either direction. **C -** Percentage of intergenic predicted enhancers linked by ChIAPPET to a nearby TSS. In the YPaxis we plot in log2-scale the fraction of regions with ChIAPPET links to a nearby TSS, for activated, silenced, random activated and random silenced enhancers. TSS – enhancer pairs are considered when all elements are located at least 3 kilobases away and as far as 100 kilobases.

**Supplementary Figure 10.**
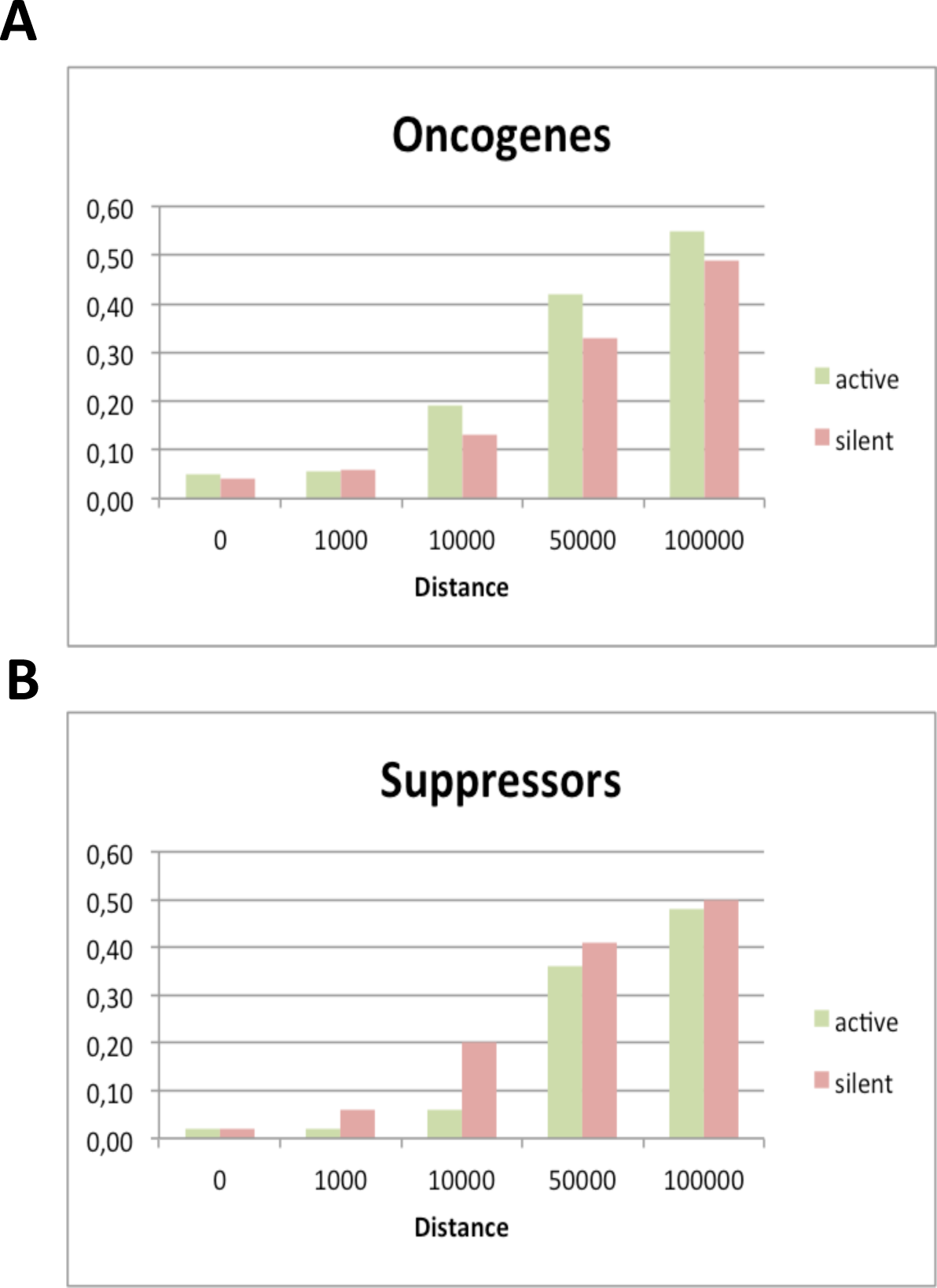
These plots show on the y-axis the proportion of the Oncogenes **(A)** or Suppressors **(B)** with an active or silent enhancer at a distance indicated on the x-axis in bp.

**Supplementary Figure 11.**
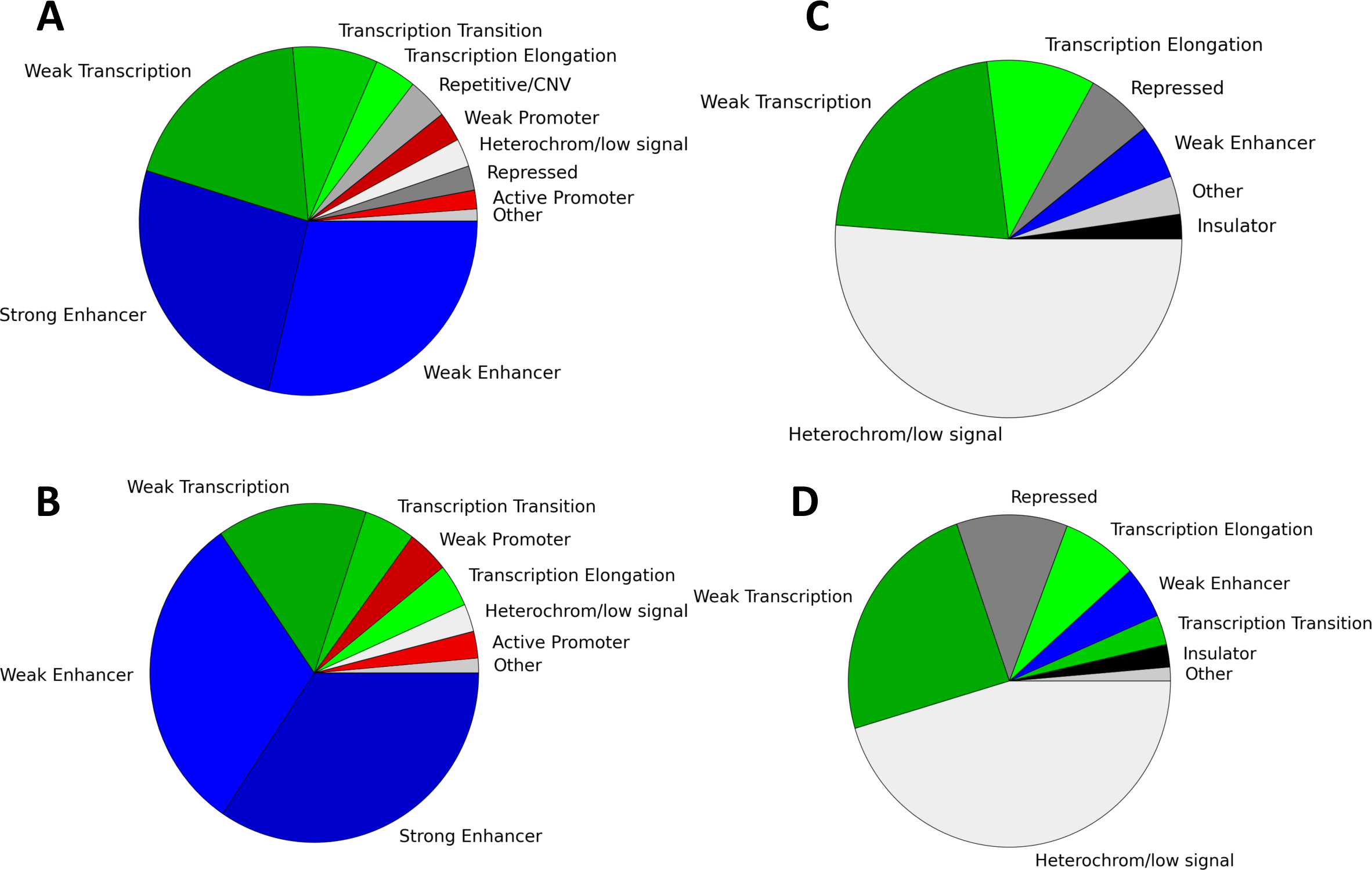
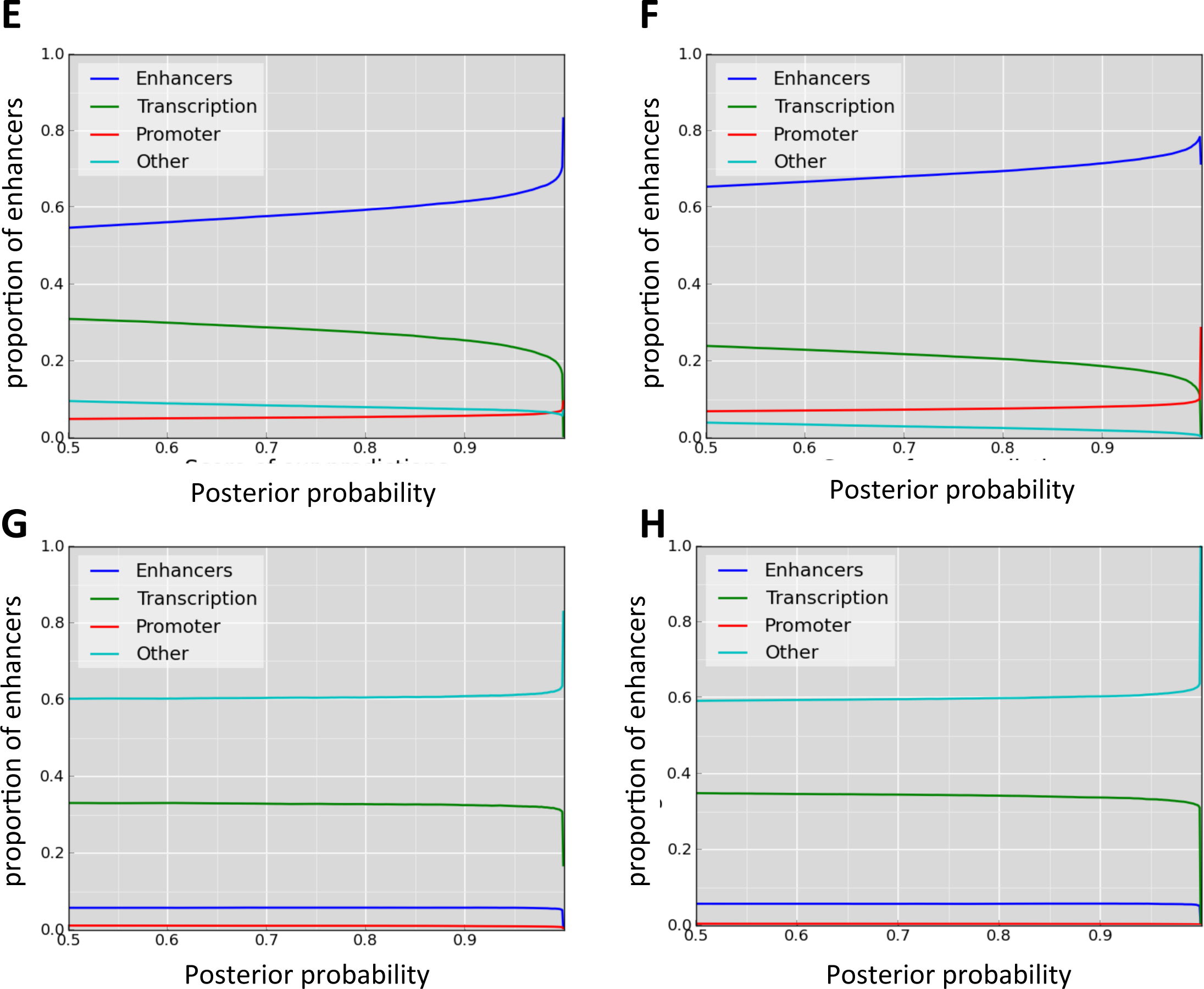
Pie charts showing the overlap of **(A)** active intragenic enhancers in K562 with ChromHMM classes for K562, **(B)** active intragenic enhancers in GM12878 with ChromHMM classes for GM12878, **(C)** silent intragenic enhancers in K562 with ChromHMM classes for K562 and **(D)** silent intragenic enhancers in GM12878 with ChromHMM classes for GM12878. Proportion of overlap of predicted enhancers with ChromHMM regions as a function of the posterior probabilities, for active enhancers in K562 **(E)** and GM12878 **(F)** and of silent enhancers in K562 **(G)** and GM12878 **(H),** with regions predicted by ChromHMM in the corresponding cell line. Color codes as in supplementary figure 5.

**Supplementary Figure 12.**
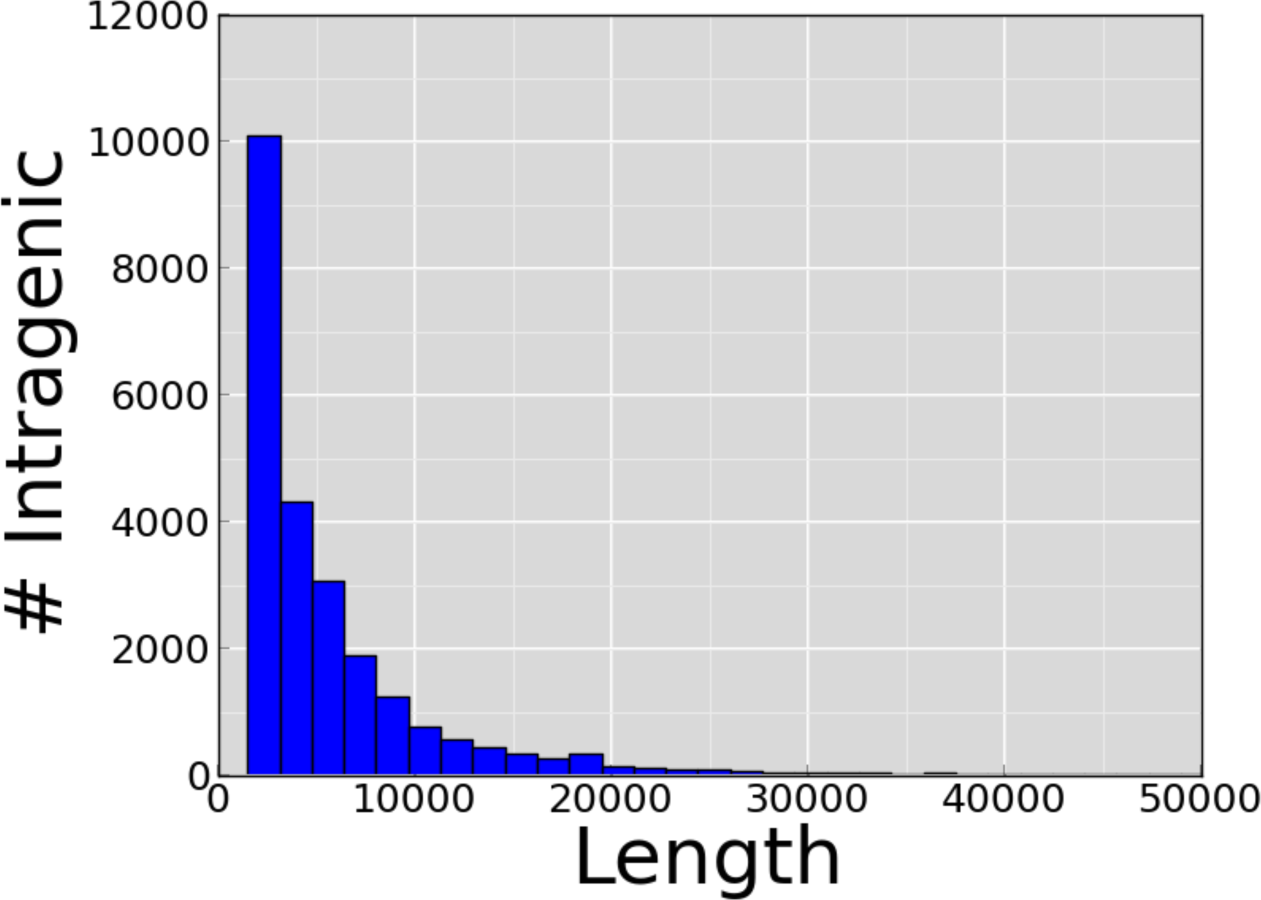
Length distribution of intragenic enhancers

**Supplementary Figure 13.**
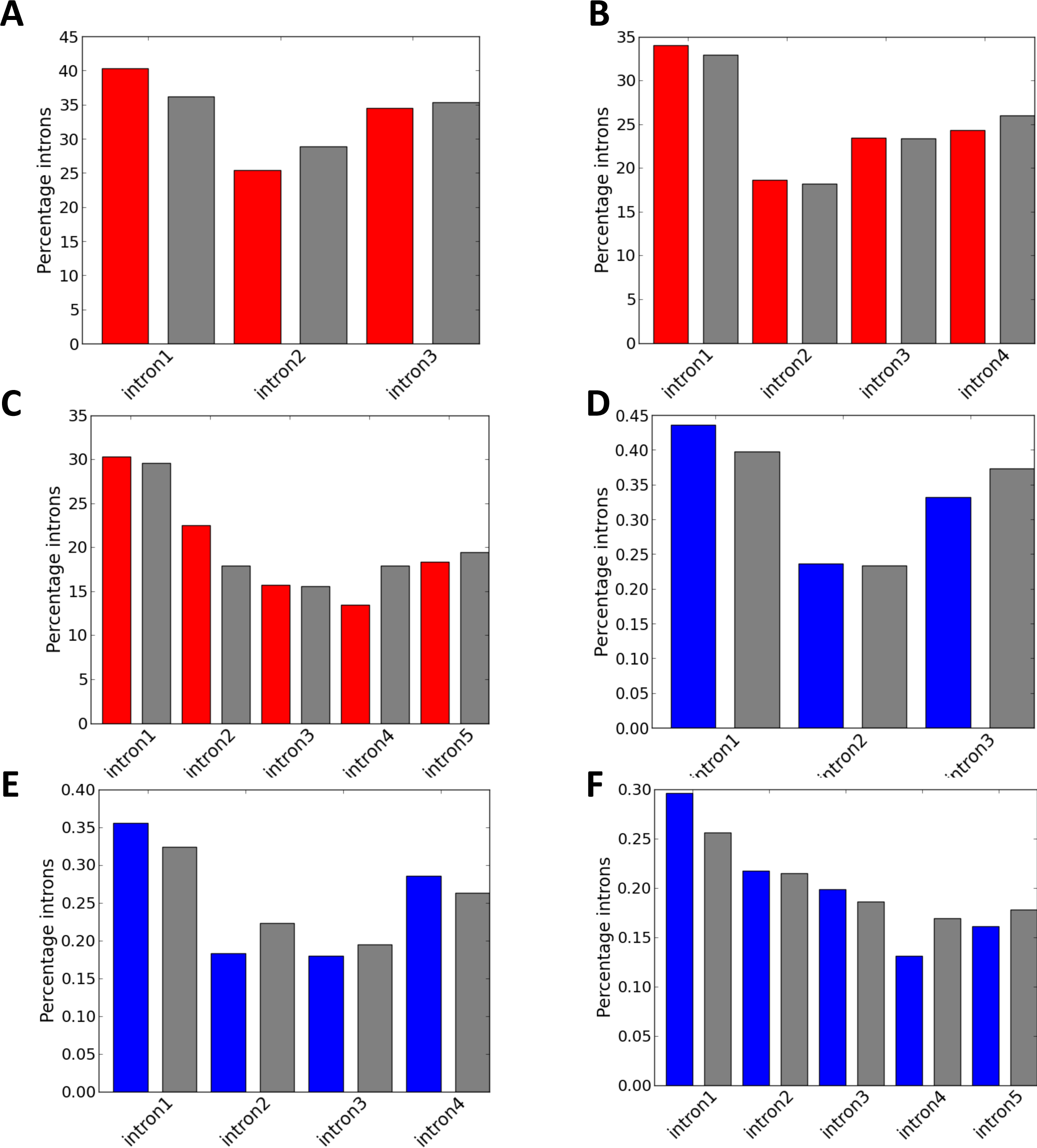
These barplots show the positional bias of intronic active and silent enhancers. The yPaxes indicate the percentage of enhancer in each intron for active enhancers (red) and randomized enhancers (gray) in genes with just 3 **(A)**, 4 **(B)** or 5 **(C)** introns. Similarly, we show the proportion in each intron of silent enhancers (blue) and randomized enhancers (gray) for genes with just 3 **(D)**, 4 **(E)** or 5 **(F)** introns.

**Supplementary Figure 14.**
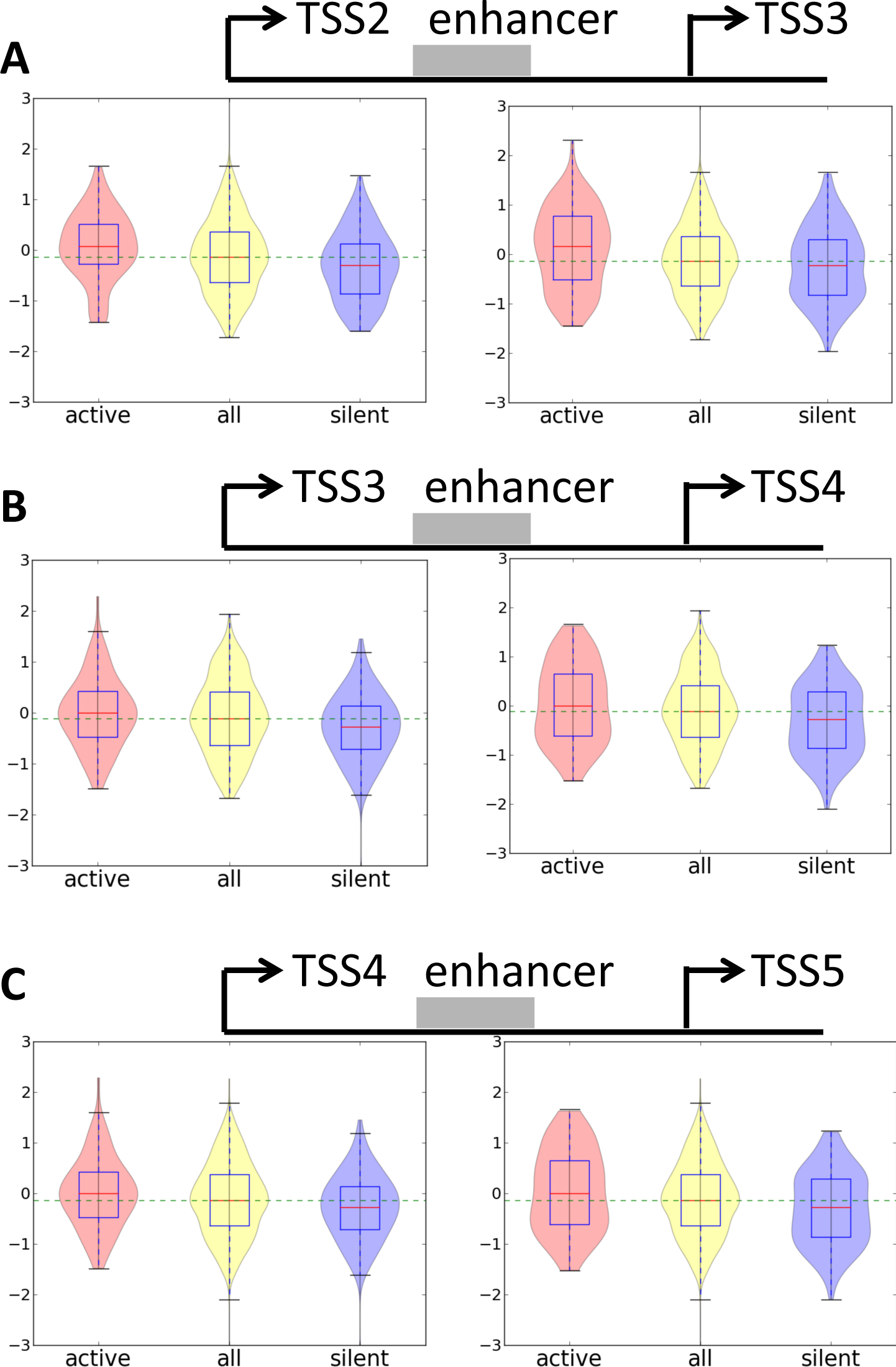
RNAPII relative levels at the TSSs fianking intragenic enhancers, for the second and third TSS (A), third and fourth and fourth and fith (C) when they fiank an active enhancer (light red), a silent enhancer (light purple) and for the whole set of putative enhancer regions.

**Supplementary Figure 15.**
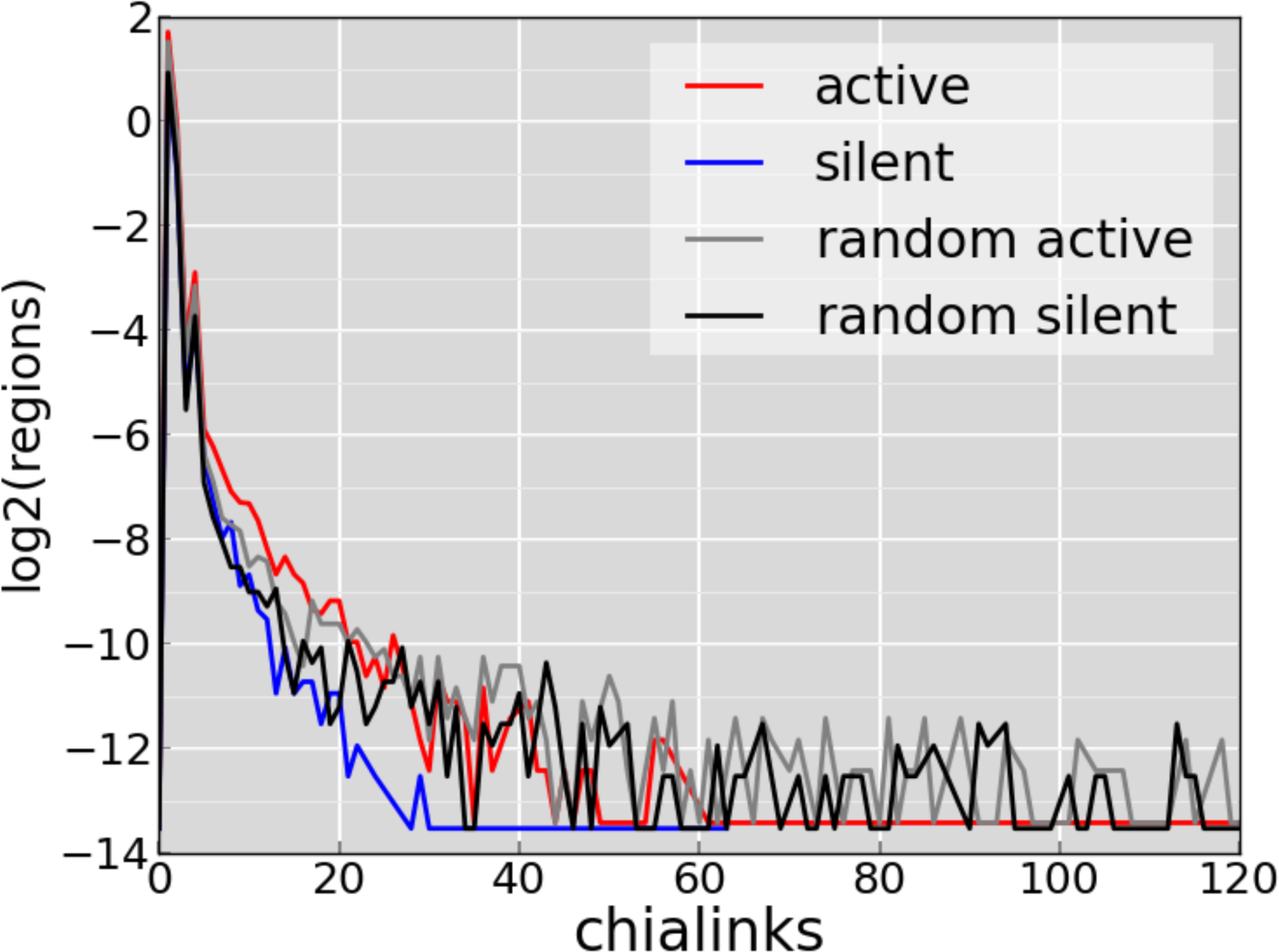
ChIA-PET links for intragenic enhancers. In the YP axis we plot in log2Pscale the fraction of regions with ChIA-PET links to a nearby TSS, for active (red), silent (blue), random active (light grey) and random silent (dark grey) enhancers. TSS – enhancer pairs are considered when all elements are located at least 3 kilobases away and as far as 100 kilobases.

**Supplementary Table 1.** First introns are longer on average. We calculated the average of every intron separating genes in the GENCODE.V7 annotation by number of introns, from 2 to 7 introns. The only case where the first intron doesn’t seem to be longer in the 2 introns genes group.

|  |
| --- |
| **Total genes with 2 intron(s): 5392** |
| Intron 1 avg. length: 7263.61183234 |
| Intron 2 avg. length: 7199.96735905 |
| **Total genes with 3 intron(s): 3089** |
| Intron 1 avg. length: 7832.92651343 |
| Intron 2 avg. length: 4888.83360311 |
| Intron 3 avg. length: 6798.17578504 |
| **Total genes with 4 intron(s): 2296** |
| Intron 1 avg. length: 7814.87108014 |
| Intron 2 avg. length: 4712.29790941 |
| Intron 3 avg. length: 4521.63719512 |
| Intron 4 avg. length: 6246.63240418 |
| **Total genes with 5 intron(s): 1863** |
| Intron 1 avg. length: 6881.82930757 |
| Intron 2 avg. length: 5561.91250671 |
| Intron 3 avg. length: 4552.50187869 |
| Intron 4 avg. length: 4049.90982287 |
| Intron 5 avg. length: 5628.71604938 |
| **Total genes with 6 intron(s): 1554** |
| Intron 1 avg. length: 6973.46718147 |
| Intron 2 avg. length: 5309.77284427 |
| Intron 3 avg. length: 4375.55984556 |
| Intron 4 avg. length: 4445.78635779 |
| Intron 5 avg. length: 4006.58687259 |
| Intron 6 avg. length: 4861.94144144 |
| **Total genes with 7 intron(s): 1290** |
| Intron 1 avg. length: 6926.83333333 |
| Intron 2 avg. length: 5174.78914729 |
| Intron 3 avg. length: 4665.16744186 |
| Intron 4 avg. length: 4541.41782946 |
| Intron 5 avg. length: 3769.95968992 |
| Intron 6 avg. length: 3817.72868217 |
| Intron 7 avg. length: 4435.38294574 |

**Supplementary Table 2.** Number of genes with and without enhancers active in either cell line that have at least one alternative splicing event that changes between the two cell lines. The regulated events were calculated as described in Methods. Events regulated according to either replicate comparison were considered for the comparison. All genes were multi-exonic.

| RNA-Seq | All genes | Genes with regulated events (cytosolic) | Genes with regulated events (nuclear) |
| --- | --- | --- | --- |
| Genes with active enhancers in K562 | 5609 | 210 | 229 |
| Genes with no active enhancers in K562 | 26740 | 266 | 294 |
| Genes with active enhancers in GM12878 | 5905 | 231 | 254 |
| Genes with no active enhancers in GM12878 | 26444 | 245 | 269 |

**Supplementary Table 3.** Enrichment analysis of the Gene Ontology process for the genes with intragenic enhancers (active or silent in K562) that also contain regulated events (|ΔΨ| > 0.1 in any of two replicates)

| GO Term | Description | P-value | FDR q-value | Enrichment | N | B | n | b |
| --- | --- | --- | --- | --- | --- | --- | --- | --- |
| GO:0044260 | cellular macromolecule metabolic process | 2.49E-4 | 1,00E+00 | 1.31 | 5363 | 1928 | 234 | 110 |
| GO:0006351 | transcription, DNA-templated | 2.8E-4 | 1,00E+00 | 1.65 | 5363 | 653 | 234 | 47 |
| GO:0097659 | nucleic acid-templated transcription | 2.8E-4 | 9.14E-1 | 1.65 | 5363 | 653 | 234 | 47 |
| GO:0010468 | regulation of gene expression | 5.82E-4 | 1,00E+00 | 1.40 | 5363 | 1224 | 234 | 75 |
| GO:0090304 | nucleic acid metabolic process | 6.09E-4 | 1,00E+00 | 1.44 | 5363 | 1084 | 234 | 68 |
| GO:0006325 | regulation of nucleobase-containing compound metabolic process | 7.02E-4 | 1,00E+00 | 1.38 | 5363 | 1293 | 234 | 78 |
| GO:0006325 | chromatin organization | 7.64E-4 | 1,00E+00 | 2.13 | 5363 | 226 | 234 | 21 |
| GO:0016568 | chromatin modification | 7.94E-4 | 9.71E-1 | 2.17 | 5363 | 211 | 234 | 20 |

**Supplementary Table 4.** Contingency tables with the gene counts with or without active enhancers in GM12878 (left) or in K562 (right), separated according to whether they have alternative splicing events that change the PSI (|delta PSI|>0.1) or not (| delta PSI| < 0.05). The delta PSI calculation was performed twice, one for each pairing of one replicate from each cell line. Other pairings did not produce any mayor changes in the results. All odds-ratios are > 1. Fisher p-values < 0.05 are marked in green

Enhancers active in GM12878
| Cytosolic sample (rep 1) |  |  |  |
| --- | --- | --- | --- |
| multi-exonic genes |  | PSI change | PSI no-change |
|  | with enhancer | 166 | 697 |
|  | no enhancer | 169 | 867 |
| Odds-ratio |  | 1,2217 |  |
| Fisher p-value |  | 0,1027 |  |

| Cytosolic sample (rep 2) |  |  |  |
| --- | --- | --- | --- |
| multi-exonic genes |  | PSI change | PSI no-change |
|  | with enhancer | 135 | 589 |
|  | no enhancer | 155 | 807 |
| Odds-ratio |  | 1,19320 |  |
| Fisher p-value |  | 0,1923 |  |

| Nuclear sample (rep 1) |  |  |  |
| --- | --- | --- | --- |
| multi-exonic genes |  | PSI change | PSI no-change |
|  | with enhancer | 184 | 633 |
|  | no enhancer | 181 | 839 |
| Odds-ratio |  | 1,3472 |  |
| Fisher p-value |  | 0,01142 |  |

| Nuclear sample (rep 2) |  |  |  |
| --- | --- | --- | --- |
| multi-exonic genes |  | PSI change | PSI no-change |
|  | with enhancer | 185 | 586 |
|  | no enhancer | 193 | 786 |
| Odds-ratio |  | 1,2855 |  |
| Fisher p-value |  | 0,0351 |  |

Enhancers active in K562
| Cytosolic sample (rep 1) |  |  |  |
| --- | --- | --- | --- |
| multi-exonic genes |  | PSI change | PSI no-change |
|  | with enhancer | 158 | 643 |
|  | no enhancer | 177 | 921 |
| Odds-ratio |  | 1,2784 |  |
| Fisher p-value |  | 0,04432 |  |

| Cytosolic sample (rep 2) |  |  |  |
| --- | --- | --- | --- |
| multi-exonic genes |  | PSI change | PSI no-change |
|  | with enhancer | 121 | 567 |
|  | no enhancer | 169 | 829 |
| Odds-ratio |  | 1,0468 |  |
| Fisher p-value |  | 0,7429 |  |

| Nuclear sample (rep 1) |  |  |  |
| --- | --- | --- | --- |
| multi-exonic genes |  | PSI change | PSI no-change |
|  | with enhancer | 174 | 593 |
|  | no enhancer | 191 | 879 |
| Odds-ratio |  | 1,3478 |  |
| Fisher p-value |  | 0,01272 |  |

**Supplementary Table 4.** Contingency tables with the gene counts with or without active enhancers in GM12878 (left) or in K562 (right), separated according to whether they have alternative splicing events that change the PSI ( $|\Delta \text{PSI}| > 0.1$ ) or not ( $|\Delta \text{PSI}| < 0.05$ ).
| Nuclear sample (rep 2) |  |  |  |
| --- | --- | --- | --- |
| multi-exonic genes |  | PSI change | PSI no-change |
|  | with enhancer | 160 | 550 |
|  | no enhancer | 218 | 822 |
| Odds-ratio |  | 1,0969 |  |
| Fisher p-value |  | 0,4423 |  |

**Supplementary Table 5.** The same as Supplementary Table 3, but the genes considered are those that do change expression between the two cell lines (Methods). All odds-ratios are > 1. Fisher p-values < 0.05 are marked in green

Enhancers active in GM12878
| Cytosolic sample (rep 1) |  |  |  |
| --- | --- | --- | --- |
| multi-exonic<br>genes, no diff-exp |  | PSI change | PSI no-change |
|  | with enhancer | 124 | 497 |
|  | no enhancer | 109 | 642 |
| Odds-ratio |  | 1,4691 |  |
| Fisher p-value |  | 0,009215 |  |

| Cytosolic sample (rep 2) |  |  |  |
| --- | --- | --- | --- |
| multi-exonic<br>genes, no diff-exp |  | PSI change | PSI no-change |
|  | with enhancer | 97 | 432 |
|  | no enhancer | 106 | 599 |
| Odds-ratio |  | 1,2680 |  |
| Fisher p-value |  | 0,1403 |  |

| Nuclear sample (rep 1) |  |  |  |
| --- | --- | --- | --- |
| multi-exonic<br>genes, no diff-exp |  | PSI change | PSI no-change |
|  | with enhancer | 127 | 463 |
|  | no enhancer | 123 | 609 |
| Odds-ratio |  | 1,3578 |  |
| Fisher p-value |  | 0,03394 |  |

| Nuclear sample (rep 2) |  |  |  |
| --- | --- | --- | --- |
| multi-exonic<br>genes, no diff-exp |  | PSI change | PSI no-change |
|  | with enhancer | 137 | 425 |
|  | no enhancer | 126 | 558 |
| Odds-ratio |  | 1,4271 |  |
| Fisher p-value |  | 0,01196 |  |

Enhancers active in K562
| Cytosolic sample (rep 1) |  |  |  |
| --- | --- | --- | --- |
| multi-exonic<br>genes, no diff-exp |  | PSI change | PSI no-change |
|  | with enhancer | 109 | 464 |
|  | no enhancer | 124 | 675 |
| Odds-ratio |  | 1,2785 |  |
| Fisher p-value |  | 0,09378 |  |

| Cytosolic sample (rep 2) |  |  |  |
| --- | --- | --- | --- |
| multi-exonic<br>genes, no diff-exp |  | PSI change | PSI no-change |
|  | with enhancer | 93 | 420 |
|  | no enhancer | 110 | 611 |
| Odds-ratio |  | 1,22972 |  |
| Fisher p-value |  | 0,186 |  |

| Nuclear sample (rep 1) |  |  |  |
| --- | --- | --- | --- |
| multi-exonic<br>genes, no diff-exp |  | PSI change | PSI no-change |
|  | with enhancer | 124 | 419 |
|  | no enhancer | 126 | 653 |
| Odds-ratio |  | 1,5332 |  |
| Fisher p-value |  | 0,002691 |  |

| Nuclear sample (rep 2) |  |  |  |
| --- | --- | --- | --- |
| multi-exonic<br>genes, no diff-exp |  | PSI change | PSI no-change |
|  | with enhancer | 117 | 392 |
|  | no enhancer | 146 | 591 |
| Odds-ratio |  | 1,2080 |  |
| Fisher p-value |  | 0,1804 |  |

**Supplementary Figure 16.**
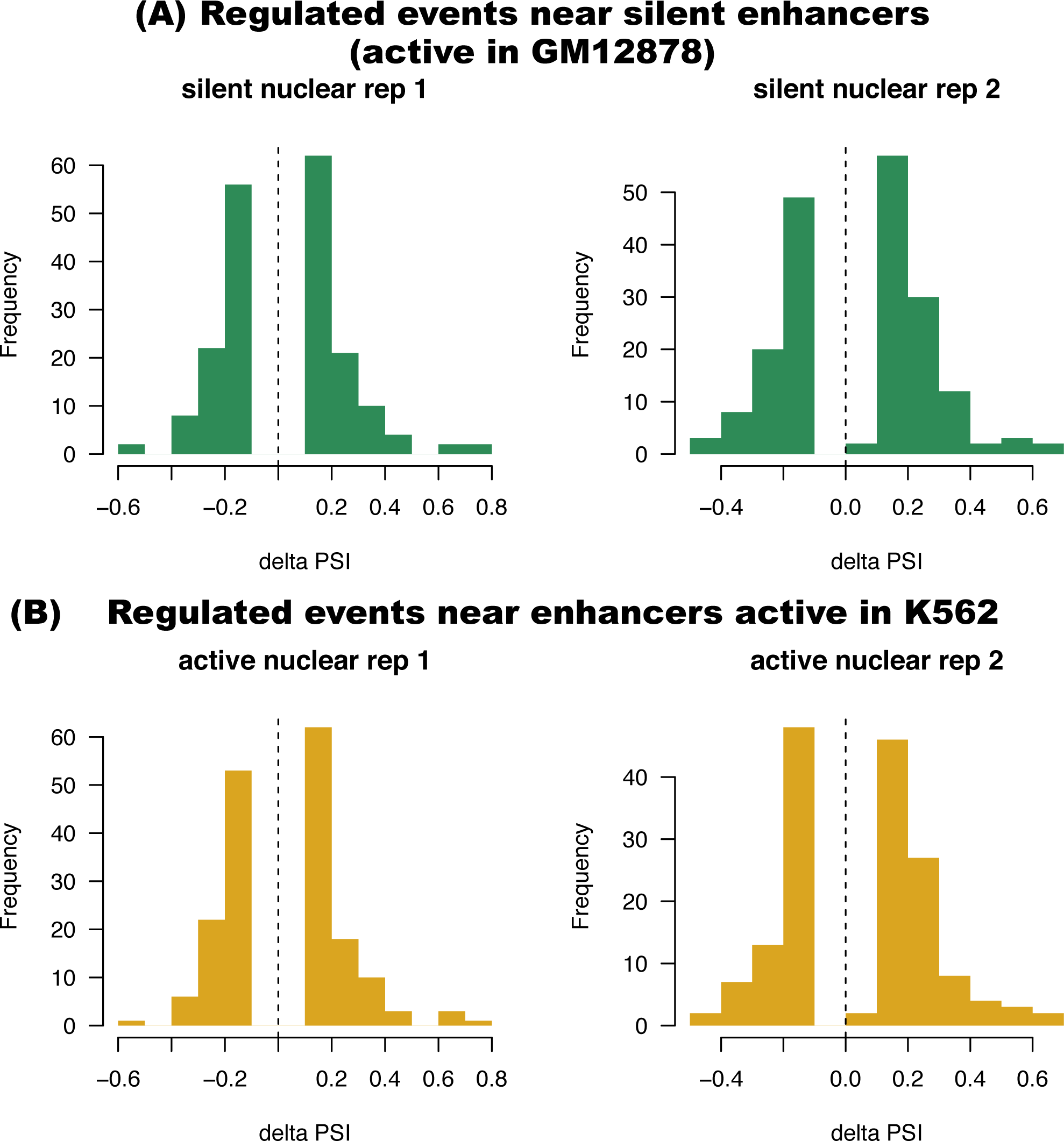
Distributions of the delta PSI values of the events regulated nearby active enhancers. Regulated events were calculated as those exon casseres that had |delta PSI| > 0.1 comparing the PSI in K562 and GM12878. Only the closest enhancer is selected for each event. Two comparisons were made, matching each replicate in one cell line with one replicate in the other cell line.

**Supplementary Figure 17.**
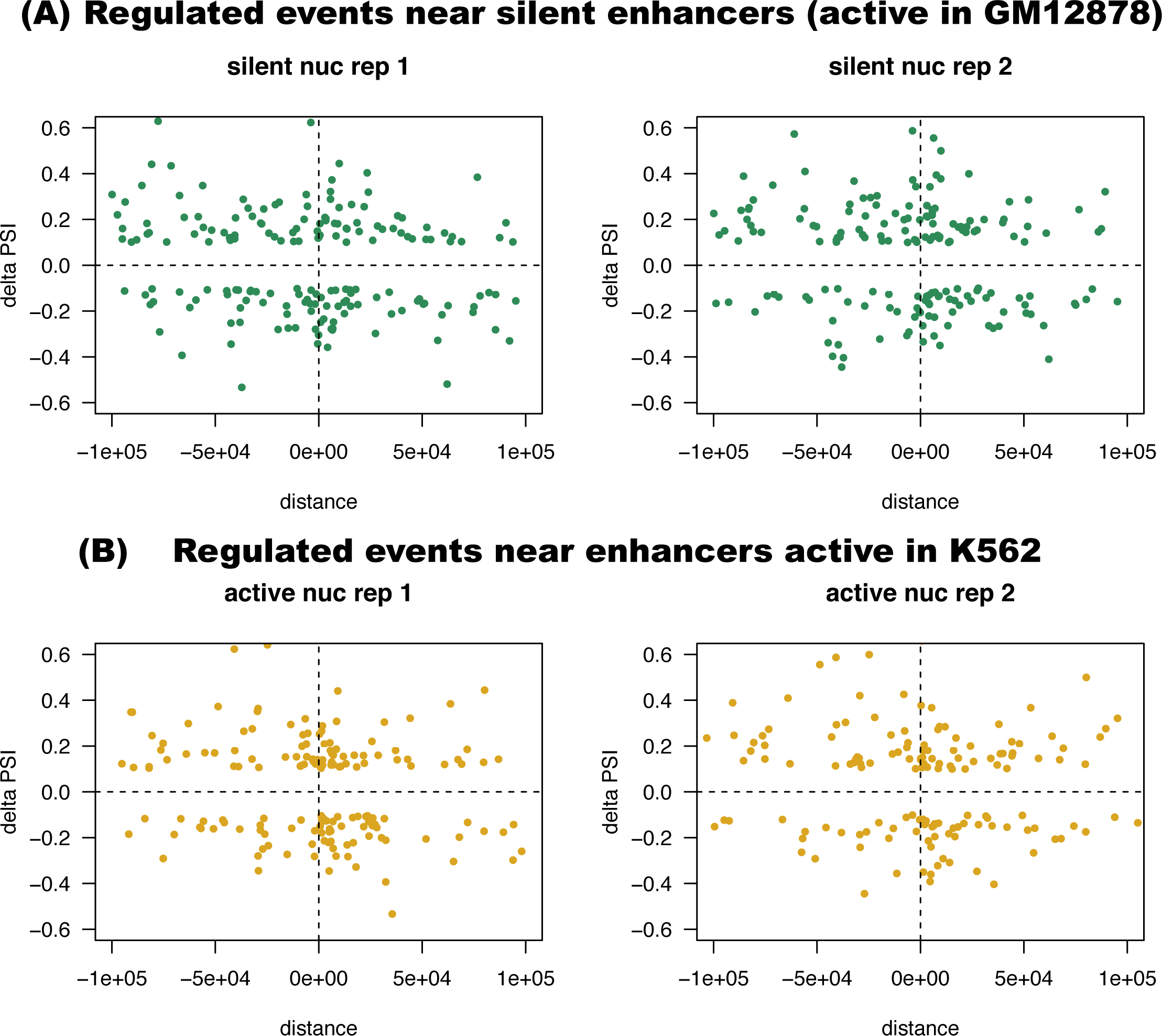
Relation between the delta PSI values (y axis), calculated as the difference of PSI between K562 and GM12878, and the distance to the closest enhancer in nucleotides (x axis). Negative and positive distances correspond to upstream and downstream positions, respectively. The distance is calculated from the middle point of the regulated exon to the middle point of the predicted enhancer. The plots show the delta PSI values calculated for the two replicate comparisons (left and right panels) for each set of enhancers, active in GM12878 (silent in K562) **(A)** or active in K562 **(B)**.

**Supplementary Figure 18.**
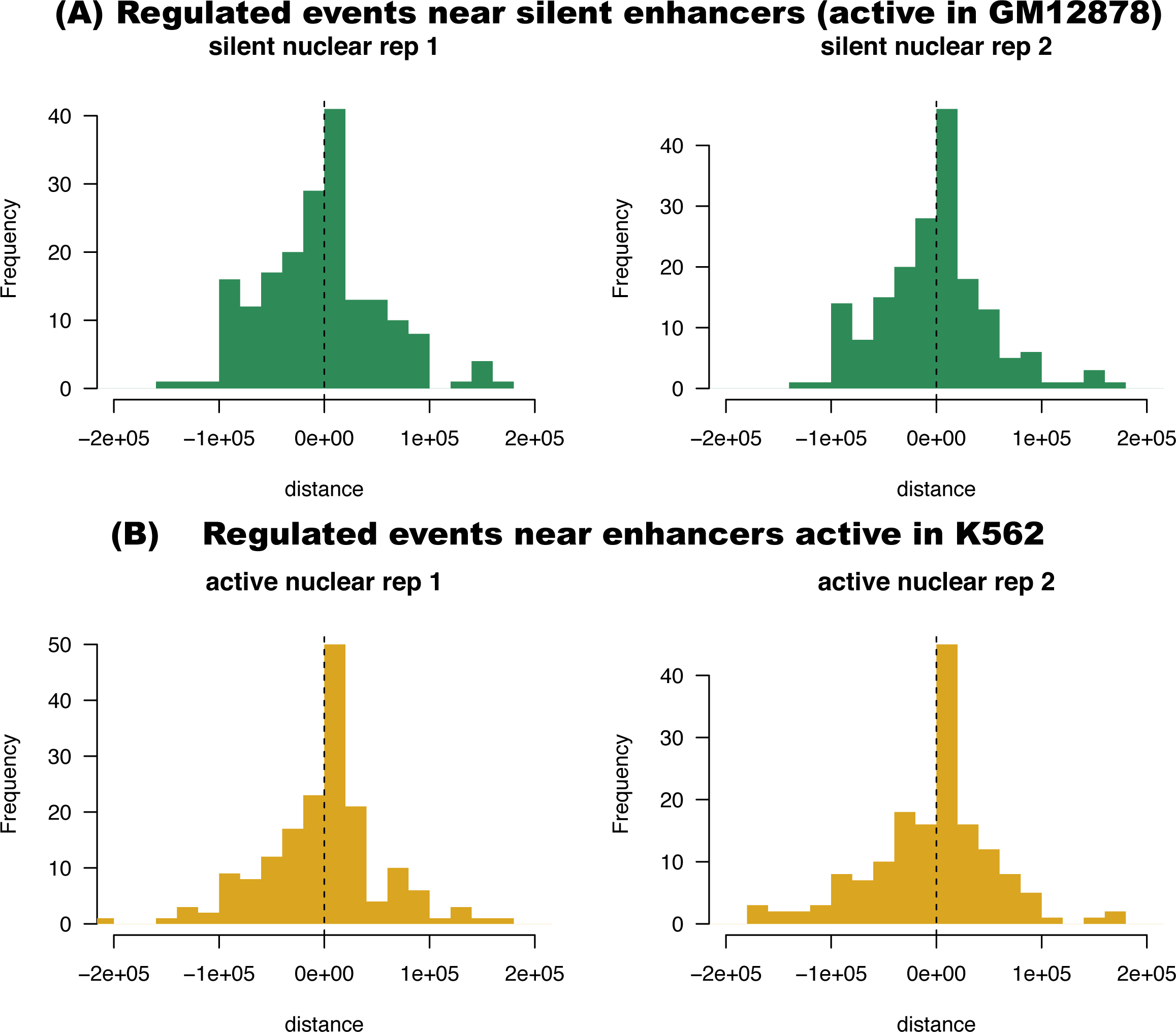
Distribution of the distances between the regulated events and the closest enhancer shown in Supplementary Figure 16. Negative and positive distances correspond to upstream and downstream positions, respectively. The distance is calculated from the middle point of the regulated exon to the middle point of the predicted enhancer. The plots show distributions for the two replicates (left and right panels) for each set of enhancers, active in GM12878 (silent in K562) **(A)** or active in K562 **(B)**.

**Supplementary Figure 19.**
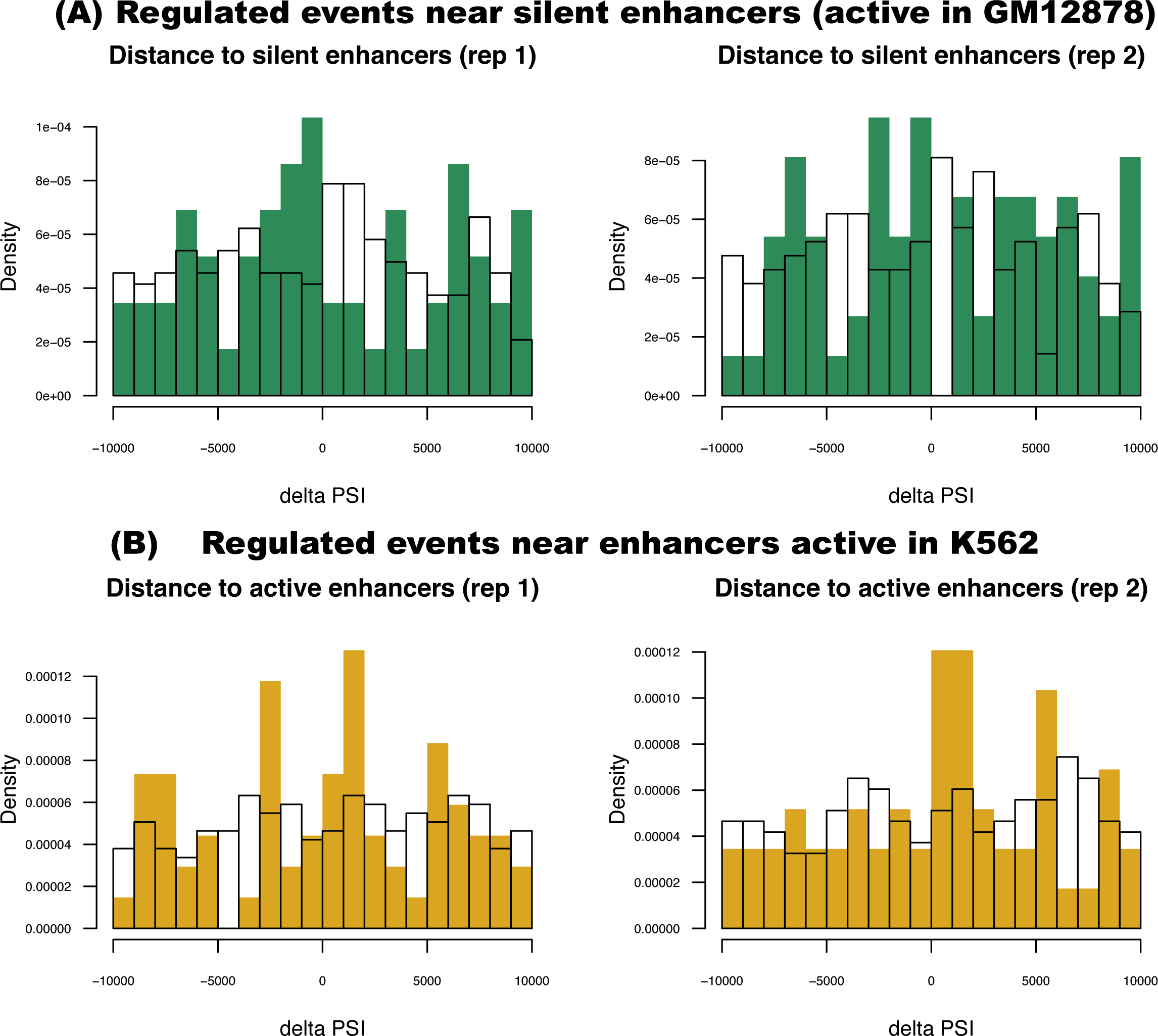
Distribution of the distances to the closest enhancer for regulated (colored bars) and non-regulated (empty bars) events. Only events genes with only regulated or non-regulated events were considered for this comparison. Negative and positive distances correspond to upstream and downstream positions, respectively. The distance is calculated from the middle point of the regulated exon to the middle point of the predicted enhancer. The plots show distributions for the two replicates (left and right panels) for each set of enhancers, active in GM12878 (silent in K562) **(A)** or active in K562 **(B)**.

